# Plagiarism in Brazil: A perspective of 25,000 PhD holders across the sciences

**DOI:** 10.1101/825026

**Authors:** Sonia MR Vasconcelos, Hatisaburo Masuda, Martha Sorenson, Francisco Prosdocimi, Marisa Palácios, Edson Watanabe, José Carlos Pinto, José Roberto Lapa e Silva, Adalberto Vieyra, André Pinto, Jesús Mena-Chalco, Mauricio Sant’Ana, Miguel Roig

## Abstract

When it comes to ownership of ideas in science, Robert K. Merton (1957) observed in *Priorities in Scientific Discovery: A Chapter in the Sociology of Science* that “what is true of physics, chemistry, astronomy, medicine and mathematics is true also of all the other scientific disciplines, not excluding the social and psychological sciences”. However, consensus over related issues, such as what constitutes plagiarism in these fields cannot be taken for granted. We conducted a comprehensive study on plagiarism views among PhD holders registered in the database of the Brazilian National Council for Scientific and Technological Development (CNPq). We collected 25,157 valid responses encompassing views and attitudes toward plagiarism from a probability sample of PhD holders across the fields, including biologists, physicists, mathematicians, and engineers as well as linguists, philosophers and anthropologists. The results suggest that core principles about plagiarism are shared among this multidisciplinary community and that they corroborate Merton’s observations. Before this study, we could only speculate that this is the case. With these data from a probability sample of Brazilian academia (PhD holders), this study offers insight into the way plagiarism is perceived across the sciences, including the literature and arts, and sheds light on the problem in the context of international collaborative research networks. The data focus on a young research system in Latin America, but, given the cultural similarities that bind most Latin-American nations, these results may be relevant to other PhD populations in the region and should provide a comparison with studies from other emerging, non-Anglophone regions.

## Introduction

Credit and priority of discovery are at the core of the scientific endeavor, and they stand out among the reasons that make plagiarism a serious ethical breach in research. As Biagioli (2012) (**1**) states, “given the crucial relationship between credit and priority in science, the plagiarism that hurts the most is one that deprives a scientist of his or her priority”. Across the globe, it seems reasonable to assume that most academics would agree that plagiarism is unacceptable and is thus research misconduct. The traditional definition promulgated by the US Office of Science and Technology Policy (OSTP) in 2000 has been a useful point of reference in many countries (**1-4**) and encompasses fabrication, falsification and plagiarism in proposing, conducting, reviewing and publishing research (**5**). However, attitudes toward plagiarism practices vary in academia, even though it is condemned as a serious ethical breach (**6,7**) - “the appropriation of another person’s ideas, processes, results, or words without giving appropriate credit” (**5**). While the definition appears straightforward at first glance, the question of which responses are appropriate, especially to plagiarism of ideas and words, does not appear to be consensual among academics (**8-10**). When it comes to scientific ideas, ownership is cultivated mostly because it is attached to the concept of priority of discovery in science, which can lead to disputes (**11**). Although notions of establishing priority are gradually changing (**12, 13**), the concept partially explains why “being the first” in science still matters today. This feeling was expressed, for example, by Francis Mojica, the Spanish scientist considered to be the first to describe the mechanism for CRISPR (clustered regularly interspaced short palindromic repeats), now a widely known gene editing tool. As Mojica recalls “… four journals said no one after the other, and the one that ended up publishing it took 6 months to get back to us…imagine you have something you know it’s big in your hands and there’s the possibility that another article that takes the originality of your work is published… I was in absolute despair”. (**14**)

Looking at scientific disputes in the history of science, Robert K. Merton described in the 1940s that “during the last three centuries in which modern science developed, numerous scientists… have engaged in… controversy” (**11**). Merton describes, for example, Galileo’ s Defense against “the Calumnies and Impostures of Baldassar Capar”, who, according to Galileo, took the invention of the “geometric and military compass” from him (**11**). A few centuries later, Lafollette (2000) conveyed this feeling of ownership among scientists: “steal my words, and you steal my authorship. Steal my idea, and you steal my identity as a scientist” (**15**). But despite the strength of claims related to ownership and authorship in the research community, plagiarism involves a cultural dimension that should not be overlooked (**16-21**). In this context, attempts have been made to explore conceptual aspects of plagiarism in the academic community, but studies are usually limited in scope (**21-24**). For the Latin American (LA) region, little is known about plagiarism *per se* in the research arena. Among the few analyses focusing on the publication system, one study that looked at the main Latin American databases SciELO and LILACS (**25**) shows that plagiarism accounts for the highest percentage of retractions: 86% of retraction notices in journals not listed by Journal Citation Reports (JCR) and 43% in JCR journals, from 2008 to 2014. These percentages are much higher than those usually found in similar studies for the Web of Science and PubMed (**26-28**). Whether this result means that Latin American editors are stricter with plagiarism than those from the US and Europe, for example, is an open question. Irrespective of this underexplored problem, LA countries such as Argentina, Chile, and Brazil have broadened their international collaborative research networks with many Anglophone and Asian countries (**29,30**). In these multicultural collaborative endeavors, shaped by a changing landscape for doing and communicating science, operating within similar research integrity frameworks is surely an asset for researchers and institutions involved (**31-34**).

One group that is clearly engaged in collaborative research and can offer valuable insights into views of and attitudes toward plagiarism is that of PhD holders. Exploring their views is timely, as they play a strong role in the production of knowledge and are most likely to lead science and scholarship in all fields, setting trends and behavior for generations to come. Exploring PhD holders’ views across the sciences is particularly relevant today, with emerging questions concerning open science and priority of discovery in an increasingly multidisciplinary landscape (**35, 12**). How to deal with these questions among newcomers in academia, for example, will surely be influenced by notions of plagiarism and originality among supervisors, who are mostly PhD holders. To our knowledge, however, this academic population has never been surveyed, in a large, quantitative study, for the way they perceive plagiarism in research. We conducted a comprehensive survey to explore these views for a probability sample of PhD holders across the sciences in Brazil.

Brazilian science has borrowed much from the US model in building its research system, strongly based on its graduate programs (**36**). This feature makes Brazil an interesting place to start investigating to what extent the conceptual framework for plagiarism in research is shared across the fields for the population of PhD holders, including biologists, physicists, mathematicians, and engineers as well as linguists, philosophers and anthropologists. Our hypothesis was that disciplinary traditions would not influence these academics’ views on core plagiarism issues in research. We present results from this national survey of PhD holders with *Curricula* recorded in the Lattes Platform, the official database hosted by the Brazilian National Council for Scientific and Technological Development (CNPq) (**37**), the main national research funding agency of the country.

## Methods

A survey addressing plagiarism, self-plagiarism and redundancy in academia was designed, as part of a project funded by CNPq. The project was approved by the Research Ethics Committee of the Institute for Collective Health of the Federal University of Rio de Janeiro (IESC/UFRJ). The survey (*SI 3*) was delivered in Portuguese, and all quotes from the literature in English were followed by a Portuguese translation, validated by two researchers who are native and/or native-like speakers of English and Portuguese. The survey was divided into five sections: A demographic section (Section I) was followed by three content sections with questions on *plagiarism* (Section II); on *self-plagiarism* (Section III); and on *redundancy* (Section IV). For most content questions, a Likert-type scale (**38**) was adopted, and for some questions in Sections II, III and IV, there was space for comments. Section V was designed for additional comments and suggestions from respondents. The survey was sent out after piloting the system and the survey form with a group of 18 Brazilian PhD holders for any necessary adjustments before widespread distribution. The delivery was conducted through an invitation by the CNPq’s Research Integrity Commission, which had established national directives for research integrity in 2011 (**39**). The invitation by the Commission was sent on October 3, 2014, to PhD holders registered in CNPq’s Lattes Platform. Established in 1999 (**37**), this Platform has been recognized as strategic for science and technology policies in Brazil. It records the profile and productivity of Brazilian academia in all fields, and its public database is “… a national standard … adopted by most of the country’s funding institutions, universities and research institutes… indispensable and compulsory for the analysis of merit and competence for funding applications…” (**37**). Information available on a researcher’s *Curriculum* includes academic background, current research areas, past and current projects, publications separated by type – papers, books, book chapters and other bibliographical contributions, such as conference abstracts. Accessing the *Curriculum* Lattes of a PhD holder affiliated with a university, it is possible to verify whether he/she is a professor, how many Master’s and PhD theses supervised are complete or underway and the identity of those who took part in the examination board. Other pieces of information usually found include peer review activity and administrative activities carried out together with teaching and research. A record of all Brazilian research groups certified by CNPq is available through the Lattes Platform – about 37,640 in 2016 (**40**) For the same year, the Platform recorded *Curricula* from 3,520,867 individuals studying and/or working in all fields of knowledge, including undergraduate and graduate students as well as Masters and PhD holders. In 2016, the latter group accounts for 6.47% of all *Curricula* in the database (**41**). The surveyed PhD holders thus made up a diverse population in terms of disciplinary backgrounds and research cultures.

The invitations to the national survey were distributed daily by CNPq’s informatics team, from October 3 to 15, 2014, to 143,431 emails, with a delivery error of about 0.2%, leading to 143,405 recipients. This number is unlikely to include more than one *Curriculum* associated with an individual, as the register in the Lattes Platform is attached to the holder’s official identification number recorded in the database. The system was open for respondents up to December 15, 2014. The inclusion criteria limited participation to individuals with a PhD whose *Curriculum* Lattes had been updated no more than six months before the survey. Invitations were made through a message by CNPq’s Commission, including access to a letter from the principal investigator. The letter described the nature and purpose of the research and included a consent form to be considered before accessing the survey form. To reduce bias and maximize anonymity, each invited respondent was randomly assigned an individual link to complete the survey; responses were then collected anonymously. Also, for this population of PhD holders, there was no restriction on inclusion of participants in terms of gender, academic status, or career experience. The emails used by CNPq were presumably valid, as they were the ones used by the Council for institutional communications with Lattes registrants. After collecting the data, we obtained a preliminary report from our web-based system designed to host the survey, and spreadsheets with the data were generated. Of the 143,405 invitations successfully delivered by CNPq, we had a return of 29,433 respondents (21%). We then established the following criteria to determine valid forms (n=25,157): responses should be from “invited users” AND from confirmed PhD holders, as requested in the survey system, AND from respondents who earned their PhD between 1950-2014 AND from PhD holders responding to at least one of the questions in Sections II, III or IV. To categorize responses by field, we used CNPq’s eight categories plus a Multidisciplinary category for “*grand* fields”: **Agricultural Sciences; Biological Sciences; Health Sciences; Exact and Earth Sciences; Human Sciences; Applied Social Sciences; Engineering; Linguistics, Language & Literature and Arts; Multidisciplinary** (http://www.cnpq.br/documents/10157/186158/TabeladeAreasdoConhecimento.pdf). For example, Biological Sciences include 13 subfields – General Biology, Genetics, Botany, Zoology, Ecology, Morphology, Physiology, Biochemistry, Biophysics, Pharmacology, Immunology, Microbiology, Parasitology, with a total of 55 specialties. Human Sciences include 10 subfields - Philosophy, Sociology, Anthropology, Archaeology, History, Geography, Psychology, Education, Political Science, Theology, with a total of 55 specialties.

Here, we show the results focusing mostly on plagiarism. The statistical significance for response patterns for all questions (**Sections II and III)** was assessed using the Kruskal-Wallis, non-parametric test. For the pairwise comparsions, the null hypothesis was rejected for p-values < 0.05 (*SI 2*) (**42**).

## Results and Discussion

Our sample source included all the target population – PhD holders with a *Curriculum* Lattes updated no more than six months before the survey, which led to 143,405 valid *Curricula*. In terms of sample size, a minimum of 14,911 participants would have been adequate for testing the statistical significance of responses, according to Equations 1 and 2 (**43**).

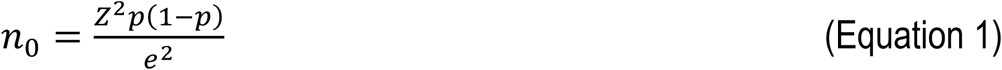

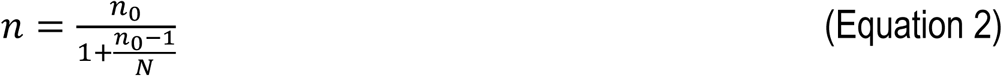

where *n*_0_ is the sample size, *Z* is the confidence level, *p* is the estimated proportion of an attribute in the population, *e* is the level of precision, *N* is the population size, and *n* is the adjusted sample size. The following values were considered for the parameters: *Z* = 2.58 (99% confidence level), *p* = 0.5 (maximum variability), *e* = 0.01 (1% acceptable sampling error), *N* = 143,405, giving an adjusted sample size of 14,911. We obtained a total of 25,157 valid respondents (68% greater than the calculated sample size). The breakdown by general field of knowledge and academic status can be seen in Table S1.

Overall, response for this national survey had a gender split of approximately 48% women and 52% men, the same as in the whole Lattes database (**41**). In terms of training, 39% of respondents stated that they had taken a post-doc, and 75% declared that they were affiliated with a public institution, such as a federal or state university or research institution. A much smaller fraction, 17%, said that they were affiliated with a private institution, and 4% with both. The remainder did not answer that question. In terms of academic positions, the three largest groups of respondents were associate professors, 55% (we made no distinction among professors categorized as “adjunto” and “associado”, as both are tenured positions in Brazil, although “associado” is for more experienced professors), researchers (non-professors), 23%, and full professors, 13% (Table S1). The “non-professors” category would include postdocs and PhD holders working on teaching and research in non-university settings or with administrative or technical jobs.

We investigated perceptions of PhD holders about plagiarism, with the definition promulgated by the US OSTP underlying most of our questions. Our respondents were distributed across the sciences, representing different disciplinary traditions, which we use in this study as a surrogate for research cultures. To qualify such a plurality of cultures, we subscribe to Whitley’s theoretical framework (2000) (**44**) where the sciences are conceived as “systems of knowledge production”. He addresses “the variety of ways in which research is organized and controlled across the sciences…” and shows “how these variations are related to different patterns of intellectual organization” (**44**). Within this framework, we assumed CNPq’s categories of “*grand* fields” as equivalent to these systems of knowledge production. We then explored views of plagiarism in research and whether these views would inform attitudes toward plagiarism in academia, particularly in real-case scenarios in the publication system. About 99% (n=24,783) of PhD holders in our sample agree (91%, n=22,898) or partially agree (8%, n=1,885) with the US OSTP inclusion of plagiarism in its definition of research misconduct (**5**) (**Q1, Section II**, Fig. S1).

On the one hand, this is not a surprising finding. As we have previously stated, PhD holders are likely to agree that fabricating, falsifying and/or plagiarizing research material would constitute research misconduct. As Biagioli (2012) (**1**) points out, “scientific credit is about content (the claim, the idea, the results, the techniques) …”. The same author adds that “the core element of scientific authorship is attribution— the professional rewards… This probably makes plagiarism the most dangerous kind of appropriation in the sciences.” (**1**). In our study, we investigated how perceptions would vary across the sciences, as plagiarism is a more controversial topic compared to fabrication and falsification (**45**). As Penders (2018) (**45**) writes, plagiarism “is often demoted to a second rank in the annals of fraud, since it is argued that plagiarism does not corrupt the content of science, only the distribution of credit, whereas fabrication and falsification do both”.

We asked respondents about the US OSTP definition of plagiarism (**Q2, Section II**), which seems clear for these PhD holders (94% agreement for **Q2a, Section II**). Our next question, in **Q3, Section II** (Fig. S2), was then targeted at the level of agreement with the definition itself, which somewhat reflects the content of other definitions in different institutions and countries (**2-4**). We found a large percentage for agreement (*Agree and Partially Agree*), a total of 98% (n=24,993) (Fig. S2). When we look at the patterns of response across the sciences for these two questions combined (**Q1 and Q3, Section II)**, a consistent pattern is found (**Fig. 1**):

**Fig. 1.**
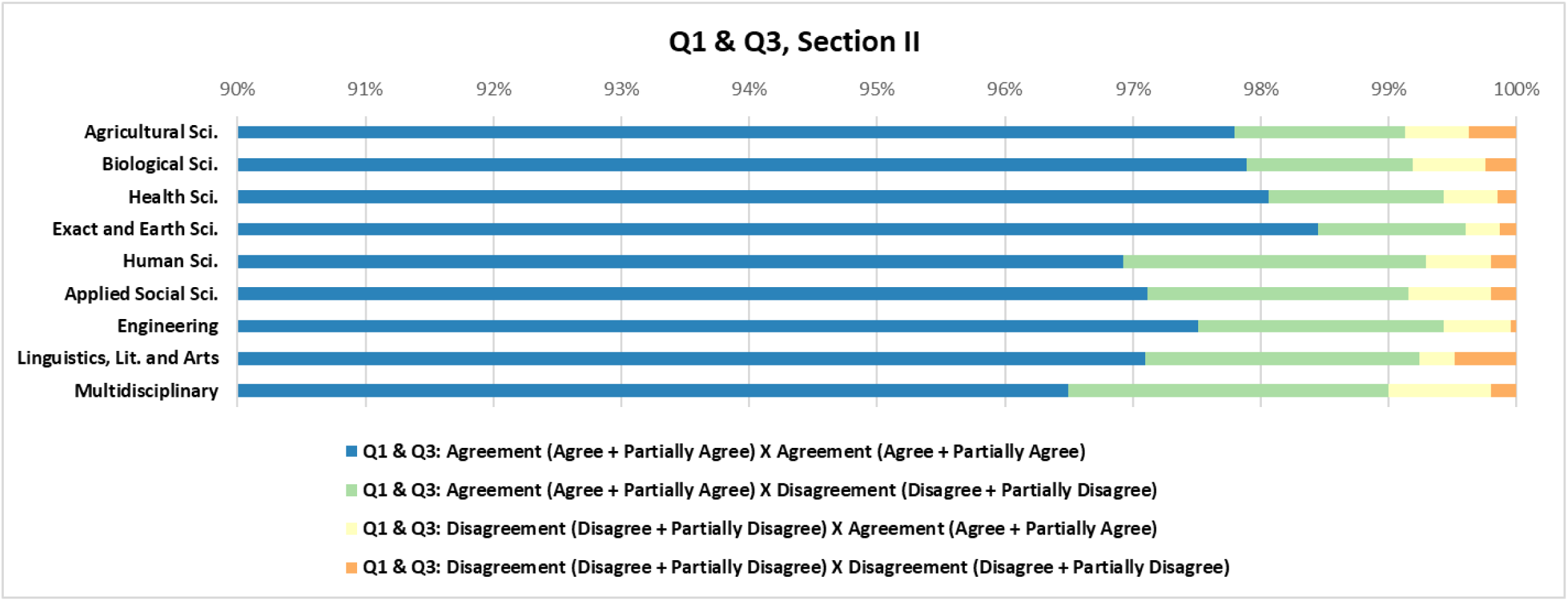
Patterns of response in all fields for respondents’ level of agreement (*Agree + Partially Agree; Disagree + Partially Disagree*) with **Q1, Section II**, on plagiarism being part of the US OSTP definition of research misconduct **& Q3, Section II** – The OSTP definition of plagiarism. The bars show the percentages of agreement and/or disagreement of respondents in each field with **both** questions.

General agreement of this population of PhD holders with the standard definitions of the US OSTP for research misconduct and for plagiarism alone reflects concerns over these ethical breaches that Brazilian academics share with the international research community. This is an important result that clarifies more than data showing that funding agencies or institutions are establishing guidelines to tackle misconduct in a given country. Consensus among PhD holders with respect to the concept of plagiarism is fundamental for an understanding of what motivates academics from such a diversity of research cultures about a sensitive issue in the production of knowledge. As expected, the US OSTP definitions are similar to those established in Brazil in 2011, by CNPq itself (**39**) and by FAPESP (São Paulo State Research Foundation) (**46**), the main state funding agency in the country. But establishment of guidelines for research integrity does not imply that academics from so many different fields would endorse the concept of plagiarism embedded in these guidelines. Given the impact that perceptions of plagiarism may have in multidisciplinary collaborative research networks, it is reasonable to ask those leading them what their views are. We thus assessed whether those patterns of response across academic fields would be consistent with the answers to a question focusing on classifying plagiarism in science (**Q4, Section II**, **Table 1**):

**Table 1.**
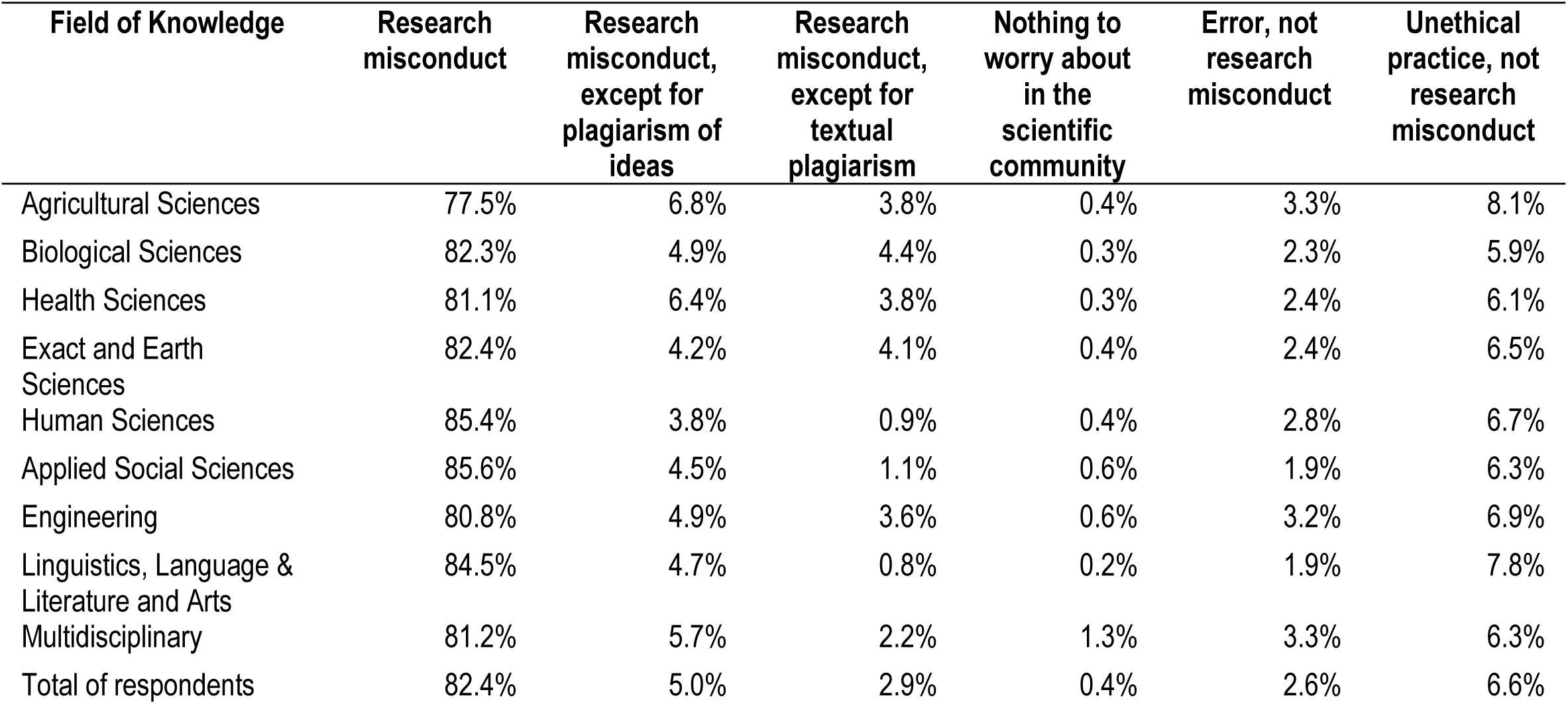
Perceptions of plagiarism in science (n=24,939) (**Q4, Section II**) **-** How do you see plagiarism in science? It is nothing that the scientific community has to worry about. It *is inappropriate, but not a form of misconduct. It is an unethical practice, but not a form of misconduct. It is a form of scientific misconduct. It is scientific misconduct, except for textual plagiarism. It is scientific misconduct, except for plagiarism of ideas*.

As shown in **Table 1**, a considerable portion (82%) of PhD holders in our sample corroborate the view that plagiarism is an instance of misconduct. This percentage is consistent with that for full agreement (91%) with the inclusion of plagiarism in the US OSTP definition of research misconduct. However, worth noting is that about 7% of respondents consider plagiarism an unethical practice and not necessarily research misconduct. This might mean that about 2,000 PhD holders in this population would be unwilling to apply severe sanctions for plagiarism. Note that percentages are quite similar across all fields. Also, one might infer that a substantial number of PhD holders in Brazil might be neglecting plagiarism practices, for example, in their tasks as supervisors. But we do not know whether this is the case. One perspective is that “the plagiarism spectrum is broad” and this practice “… seems to vary considerably in terms of seriousness, ranging from clear cases of scientific misconduct to fairly insignificant deviations from good research practice”. (**47**) This approach may also explain why almost 3% of respondents consider that plagiarism is only an error. On the one hand, one may claim that the resulting numbers represent a significant challenge to ensuring that most researchers operate within a similar research integrity framework. On the other, the result reinforces views already expressed in the research community, that “scientific plagiarism—a problem as serious as fraud—has not received all the attention it deserves” (**48**). Apart from these possible interpretations, note that only 0.4 % see plagiarism as “nothing to worry about in the research community”.

Respondents from Human Sciences, Applied Social Sciences and Linguistics, Language & Literature and Arts show the highest percentages for full agreement concerning research misconduct and plagiarism of text, with the smallest percentages (1%) for *It is scientific misconduct, except for textual plagiarism*. Agreement or partial agreement with **Q1, Section II** (Fig. S1) was prevalent for all fields, despite small differences in percentages, indicating that respondents endorsed the idea that plagiarism practices constitute misconduct. However, patterns of response for **Q1** combined with each option for **Q4, Section II** (Fig. S3) reveal some inconsistencies or lack of consensus on whether plagiarism is *misconduct, except for text or ideas*, *is an error* or *an unethical practice*.

These results are in accordance with those from a small study in Brazil indicating “lack of consensus about the appropriate limits of borrowing from the literature” among a group of scientists in biomedical science, engineering, chemistry, physics, computer science, and medicine (**22**). It is also consistent with other studies and commentaries, including one on perceptions among researchers from, say, more experimental sciences, who considered that plagiarism of text might be less severe than plagiarism of ideas and results (**19, 49**). Irrespective of these peculiarities, the response from this population of PhD holders suggests that views of core plagiarism issues are shared across research cultures. Most of our respondents thus consider plagiarism severe enough to lead to sanctions in the publication system, such as a retraction. We asked them about this issue, posing a real-case scenario (**Q5, Section II**), involving a retraction of a publication in Nature Reviews Genetics, in 2010 (**50**). The retraction was for misappropriation of ideas and text from a paragraph of a manuscript that was undergoing review (**50, 51**). The case is provocative, as the “epistemic impact” (**52**) of the retraction seems minor. But the problem has broad ethical dimensions framing the relationship among authors, reviewers and editors (**51**). After presenting the case, we first asked respondents whether they agreed that plagiarism of text would justify a retraction [in general terms] (**Q5a, Table 2**). We then asked whether they agreed with the retraction in that particular case (**Q5b,** Table S2). Most respondents agreed or partially agreed (88%) that plagiarism of text is a reason for a retraction (**Table 2**).

**Table 2.** Plagiarism and retractions (**Q5, Section II**). *Recent surveys indicate an increase in plagiarism in scientific publications. Many of the cases have led to “retractions” of scientific papers (cancellation of publications). In this context, in 2010, Nature Reviews Genetics (NRG) retracted a review article for textual plagiarism [Nature Reviews Genetics 11:308(2010]. The plagiarism involved a single paragraph that had been paraphrased from an article submitted to Plant Science. The author of the NRG review was a referee for the Plant Science paper but she failed to cite it when she wrote the NRG review. In the retraction notice the NRG editors stated that the misappropriated paragraph was plagiarized and that the author of the NRG review had presented the ideas and hypotheses found in the original paragraph as if they were her own*. ***Q5a Do you agree that plagiarism of text is a reason for a retraction?*** (n=25,029)

| Field of knowledge | Agree | Partially Agree | Disagree | Partially Disagree | I don't know |
| --- | --- | --- | --- | --- | --- |
| Agricultural Sciences | 56.7% | 30.8% | 2.6% | 5.6% | 4.3% |
| Biological Sciences | 56.8% | 30.8% | 2.7% | 6.2% | 3.5% |
| Health Sciences | 60.5% | 29.0% | 2.0% | 4.8% | 3.7% |
| Exact and Earth Sciences | 55.5% | 30.5% | 2.4% | 5.5% | 6.2% |
| Human Sciences | 68.2% | 22.1% | 1.7% | 3.3% | 4.7% |
| Applied Social Sciences | 65.0% | 24.3% | 1.7% | 3.8% | 5.2% |
| Engineering | 56.4% | 29.6% | 2.1% | 4.8% | 7.0% |
| Linguistics, Language & Literature and Arts | 67.3% | 23.6% | 1.4% | 3.1% | 4.6% |
| Multidisciplinary | 61.5% | 28.1% | 2.1% | 3.7% | 4.6% |
| Total of respondents | 60.6% | 27.8% | 2.1% | 4.7% | 4.8% |

Again, as shown in **Table 2**, Human Sciences (90%) and Applied Social Sciences (89%), as well as Linguistics, Language & Literature and Arts (91%) have the highest percentages for agreement (*Agree or Partially Agree*), together with the Health Sciences (90%) and the Multidisciplinary field (90%). These percentages are likely to reflect the stronger adherence to written arguments and, in many cases, authorial voice in many works in these fields, with a marked presence of qualitative research. This may explain, for example, why the Health Sciences (90%) yielded a similar percentage. It includes subfields such as nursing, public health, and nutrition, in which qualitative research has a strong role. Note that authors in Engineering (7%) and Exact and Earth Sciences (6%) have the highest percentages for “I don’t know”. One factor may be that the Exact and Earth Sciences have a different relationship with the text, compared to authors in the so-called “soft sciences”.

Bouville (2008) (**53**) states that “an experimental result that is described using different words is not a different result and its scientific importance is not affected by the wording… the core of science are facts and theories, not words”. Thus, doubts among respondents in these more “hard-science” fields are not a surprise. The same group expressed similar doubts when asked whether the case in **Q5b, Section II** (Table S2) should lead to a retraction: “I don’t know” received the highest percentages for Engineering (12%) and Exact and Earth Sciences (11%). The level of agreement (*Agree or Partially Agree*) for this question had similar percentages – above 81%, except for the Exact and Earth Sciences (79%) and Engineering (79%), which were almost identical. The higher percentages for “I don’t know” for these fields were followed by 10% for Multidisciplinary, 10% for Human Sciences, 9% for Applied Social Sciences and 9% for Linguistics, Language & Literature and Arts (Statistical significance for responses to these and other questions are shown in *SI 2*).

This real-case scenario is not simple, as it involves a delicate situation of misappropriation of research material in the context of peer review, but the patterns of response somewhat reflect a current trend in the publication arena: discussions on the correction of the literature and publication ethics are still led by the bio and health sciences (**26-28, 54-56**). However, irrespective of this imbalance in the publication context, a question brought up by **Q4**, **Q5a** and **Q5b, Section II** is that attitudes toward plagiarism of text for PhD holders associated with the Health Sciences, Human Sciences and Social Sciences, and with Linguistics, Language & Literature and Arts would be stricter when it comes to retractions for textual plagiarism. For the Human Sciences, Linguistics, Language & Literature and Arts, these results are consistent with previous analyses, as shown by Horbach and Halffman (2019) (**57**). We looked at patterns of response for this case (**Q5a** and **Q5b, Section II**), considering years since earning the PhD - **Group1**: up to 5 years; **Group 2**: 5-10 years; **Group 3**: 10-15 years; **Group 4**: 15-20 years; **Group 5**: 20-25 years; **Group 6**: more than 25 years. Overall, patterns of response are quite similar but suggest that a longer career tends to make respondents stricter about this problem (Fig. S4 and S5).

We also investigated respondents’ agreement with the US OSTP definition of plagiarism (**Q3, Section II**) vis à vis their attitudes toward another real-case scenario in the publication system involving self-plagiarism (**Q2, Section III**) (**Fig. 2**). Patterns of response in **Fig. 2** suggest that for respondents across all fields, self-plagiarism is not easy to define. The patterns are more varied, compared to those for **Q3, Section II** (Further details in Fig.S2, *SI 1*).

**Fig. 2.**
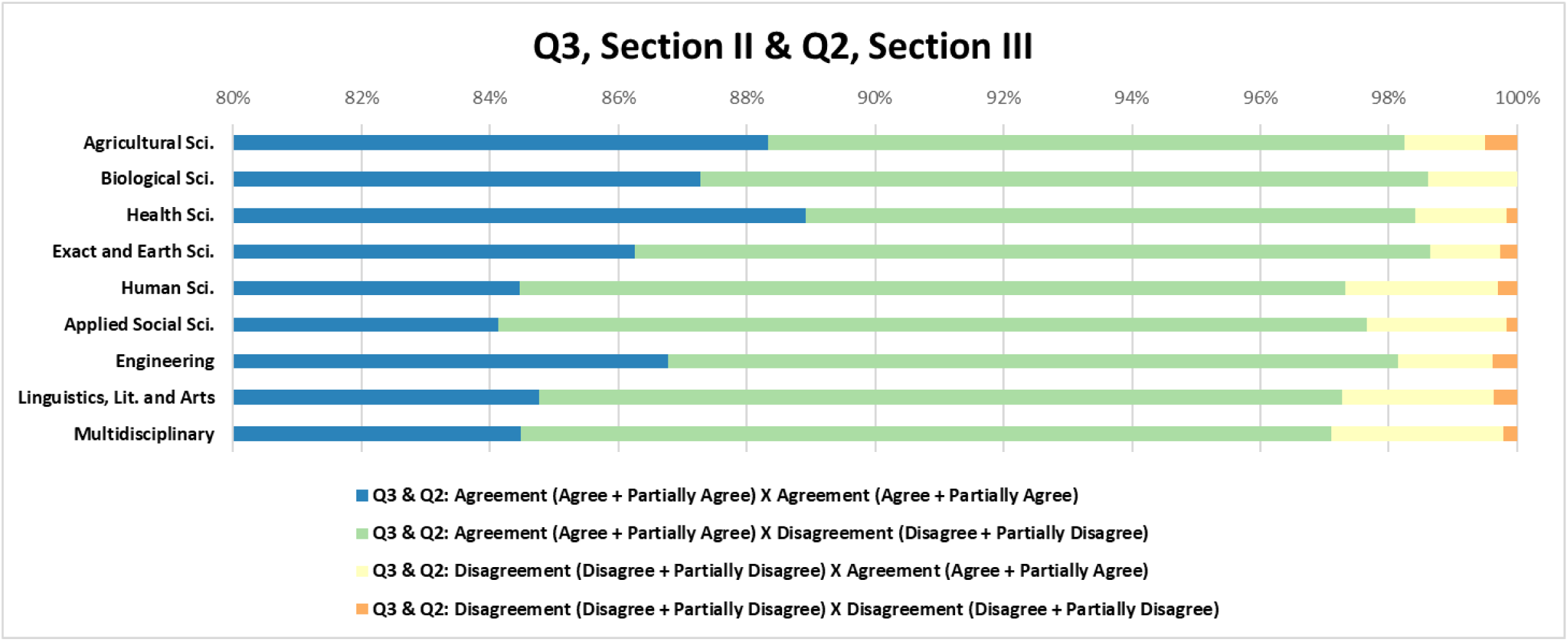
Patterns of response across all fields for respondents’ level of agreement (*Agree + Partially Agree; Disagree + Partially Disagree*) with US OSTP’s definition of plagiarism (**Q3, Section II**) (n=24,993) and with (**Q2**, **Section III**) (24,430) *Cases of self-plagiarism in science have been claimed using Crosscheck. However, there is little consensus as to how much material reused by an author, borrowing from his own publication, would be self-plagiarism An author recently accused of self-plagiarizing from one of his previously published articles claimed: “I cannot plagiarize myself; those words are mine.”* (When is self-plagiarism ok? The Scientist, Sep 2010). *Do you agree with this view? 1– Agree; 2 – Partially agree; 3 – I don’t know; 4 – Partially disagree; 5 – Disagree*. The blue bars show the percentages of respondents in each field who agreed or partially agreed with **both** questions.

As Samuelson (1994) (**58**) states, “self-plagiarism is sometimes unlawful and unethical. Other times it is unethical but not unlawful”. In Brazil, for example, individuals can reuse their authorial works, according to the country’s legislation; we could speculate that this legal framework may influence the views of the Brazilian academia about self-plagiarism. Nevertheless, a reasonable reading is that the higher percentages for “I don’t know” for **Q2, Section III,** across all fields, corroborate the idea that self-plagiarism may be considered as a questionable research practice but not as research misconduct (**58,59**).

The publication system has cultivated the notion of self-plagiarism as, for example, “the use of part of one’s own previously copyrighted work, such as sections of a review of the literature, or descriptions of design and methods sections in a research study” (**59**). Although self-plagiarism, including text reuse, goes beyond copyrighted work and is a concern in the publication system (**58-60**), it has far less ethical weight in academia when compared to plagiarism (**61,62**). Overall, these patterns of response for self-plagiarism corroborate a lack of consensus about the limits of reusing one’s own words and ideas in the context of scholarly production (**57**). Self-plagiarism will probably remain a controversial issue in academia. However, addressing self-plagiarism and plagiarism among early-career researchers is timely, and they should be invited to have a say in the conversations. They have increasingly been challenged with demands posed by an open science culture that is reshaping the concept of priority of discovery and the reward systems of science. Accordingly, we asked the PhD holders about plagiarism among graduate students. As we have pointed out, Brazilian science is mostly based on its system of graduate programs, which makes this and other related questions essential for this study. Their responses suggest that the incidence would be relatively low, although, at the policy level, using plagiarism detection software at Brazilian universities is not a common practice. Concerning the use of plagiarism detection software in the publication system, 37% to 48% of respondents indicated that they were not aware of these tools (**Q1a, Section III,** Fig. S6). These factors make it difficult for respondents to estimate the frequency with which they come across plagiarism in their tasks as, for instance, members of examination boards for evaluation of theses. When asked (**Q7, Section II**) *Have you ever come across any case of plagiarism (partial or total) by any graduate student (not necessarily from your Program) in the last four years*?, percentages for “No” varied from 50% to 65% for Master’s and from 66% to 81% for PhD theses (**Q7a, Section II,** Tables S3 and S4, respectively). One caveat in this result is that only respondents associated with graduate programs would have the experience necessary for an informed response. As about 23% of respondents were non-professors, it is likely that most of them were not acting as thesis supervisors. Aiming to broaden our understanding of the possible dimensions of plagiarism in the context of graduate studies, we set out the following condition: *In the context of plagiarism, it should be highlighted that for Brazil* (**Q8, Section II**), *Graduate students are not familiar with the international concept of academic plagiarism* (**Q8a, Section II**). Interestingly, about 72% of respondents agreed or partially agreed with the statement. However, about 11% declared “I don’t know” (Fig. S7 and S8 - Further details in *SI 1*).

### Conclusions

The results of this study broaden the research community’s perspective on the way plagiarism is perceived among PhD holders, who play an essential role in the production of knowledge and in setting trends in behavior in academia. Overall, for most questions in the survey, the level of agreement among this broad sample of academics in Brazil seems to corroborate Merton’s (1957) observation for ownership of ideas among academics - “what is true of physics, chemistry, astronomy, medicine and mathematics is true also of all the other scientific disciplines, not excluding the social and psychological sciences” (**11**). Before this study, academia could only speculate that this is the case.

Yet, there appear to be peculiarities in how these definitions are applied to the publication system, which are supported by significant differences in response patterns, according to Kruskal-Wallis analyses (*SI 2*). Although PhD holders from all sciences agree that plagiarism is an instance of research misconduct, their attitudes toward the type of plagiarism seem to be influenced by disciplinary cultures, which, as stated earlier, we use as a surrogate for research cultures. In this study, we found that PhD holders in the Human Sciences and Social Sciences, for example, have a more stringent view of misattribution of words and ideas in research.

From a broader perspective, PhD holders’ views that corroborate the idea that plagiarism is only an error but not misconduct may be propagating misperceptions about the problem that persist from their early grade-school years (**63,64**). One concern is perpetuating these views in mentor/mentee relationships. Also, the percentages of “I don’t know” for some survey questions are worth noting. Even if they are comparatively small or regarded as irrelevant, these uncertainties should be seen, particularly for Brazil, as a red flag worthy of further investigation.

Our results shed light on the way plagiarism is perceived in a young research system such as that of Brazil, whose academic structure, particularly its system of graduate programs, has been historically influenced by that of the US. These programs are the source of most of the science produced in Brazil in all fields. By drawing upon the US OSTP definitions, we show the extent to which they have been shared among young and experienced scholars in a developing country that contributes to multiple collaborative research networks.

Finally, this study offers a perspective on the way plagiarism is perceived across the sciences, including the literature and arts, in a young research system in Latin America. However, given the cultural similarities that bind most Latin-American nations, results may be relevant to other PhD populations in the region and should provide a comparison with studies from other emerging, non-Anglophone regions. In a 2012 publication, some of us reported on the launch of this national survey (**65**), aiming “to record trends among different disciplines, and to identify culturally sensitive factors that might serve to inform national, and possibly international, RI/RCR training programmes for young researchers.” We believe these results are an asset to the planning of such policies, which should combine the efforts of PhD holders across the fields.

### Study Limitations

Among the caveats in the interpretation of these results is the fact that a fraction of these PhD holders are not active in research, as they could be working for administrative or technical sectors (**41**). On the other hand, there is, or should be, an expectation that at this level of training individuals have a basic understanding of fundamental aspects of research integrity, including matters of plagiarism. After all, earning a doctoral degree usually entails some exposure to practices involved in producing knowledge in a given disciplinary field, interacting with the related literature, with supervisors, and with a research culture. For the graduate system in Brazil, this type of exposure and interaction would be expected. We do not have data on the fraction of PhD holders in our sample who have acted as authors and reviewers for international journals. However, as about 75% of our respondents are professors, and given the way academia is assessed in Brazil, it is reasonable to assume that a considerable fraction of them are active in the publication system. For most universities and research institutes in the country, publication in international journals is an asset for career promotion - especially for the natural sciences, health sciences and engineering, but not restricted to these fields. These sciences make up the largest portion of our respondents (64%). It is important to note that we included in our survey only PhD holders who had updated their CVs in the last six months prior to the survey. This criterion is relevant for this type of study as almost all funding agencies in Brazil require that the CV Lattes be updated shortly before submitting a grant application. This criterion probably biases the study towards respondents who are more active in academia.

Another caveat concerns the survey instrument, which uses a Likert-type scale for most of the questions. One issue is the scale direction effect – endorsement may be biased for scales that provide the first option in positive (*Agree or Partially Agree*) or negative mode (*Disagree or Partially Disagree*). This endorsement bias would lead to the so-called primacy effect, *i.e.*, the tendency for choosing the option provided first, irrespective of being positive or negative (**64**). However, as Liu and Keusch (2017) (**67**) have shown, “empirical studies report mixed findings with regard to the influence of the direction of rating scales on response”. The same authors conducted an experimental study to assess this effect and noted that “the scale direction does not impose a significant influence on the substantive content latent class variables, with or without controlling for response styles” (**67**). As for social desirability bias or acquiescence bias (the tendency for choosing a positive response regardless of order) (**68,69**), we believe we have been able to minimize this type of bias, varying the way we address plagiarism concepts and contexts in the survey instrument. Also, the instrument does not include potentially embarrassing questions, such as asking these research participants about their own behaviors concerning research misconduct. This element should reduce, for example, the effects of social desirability bias (**68,69**). Additionally, we surveyed a diverse population across the sciences, across Brazilian states and institutions – with respondents from diverse backgrounds, including mathematicians and engineers as well as linguists, philosophers and biologists, different professional/career experience, and different research cultures experienced during their PhD studies. Any effect driven by scale response order, acquiescence or social desirability bias should not be big enough to invalidate these results. It thus seems unlikely that our results are offering an unreliable picture of the problem studied. Nevertheless, additional large studies are planned to deepen our understanding of variables influencing the way researchers and educators deal with plagiarism notions and practices in different countries.

## Acknowledgments

We thank Professor Jacqueline Leta at the Science Education Program of IBqM/UFRJ for her contribution to the initial stages of the survey design. We are especially indebted to CNPq (grant 486220), mainly to Professor Paulo Sérgio Beirão, at the Federal University of Minas Gerais (UFMG), for his support during the whole process of implementing the national survey in Brazil. Professor Beirão helped establish the Research Integrity Commission of CNPq in 2011, which he chaired until 2015. It was in his capacity as chair of this Commission that he collaborated to involve many Brazilian PhD holders in this endeavor. At CNPq, we also thank Alerino dos Reis e Silva Filho for his assistance with practical matters related to the operationalization of the survey. We also thank the informatics team at CNPq. Professor Antonio de Figueiredo is also acknowledged. He discussed with some of us possibilities for developing a custom system for the survey, setting up a team to work on the system at Scire/COPPE/UFRJ. Professor Nick Steneck, at the University of Michigan, is also acknowledged for his contribution through early discussions on the project and scope of the study. Our thanks also go to Gabriel Elias, a former undergraduate student in biomedicine at UFRJ, who, during his training in science communication in the first author’s group contributed to solving some operational issues in the dataset, particularly those related to the *corpus* generated for qualitative analysis. Professor Carlos André Pérez, at the Federal Institute of Education, Science and Technology of Rio de Janeiro (IFRJ), is acknowledged for his critical reading of this manuscript and relevant comments.

## Supporting Information

**Other supplementary materials for this manuscript include the following:**

Kruskal-Wallis statistical test (separate file), *Supporting Information (SI 2)*

Survey Instrument (separate file), *Supporting Information (SI 3)*

**Fig. S1.**
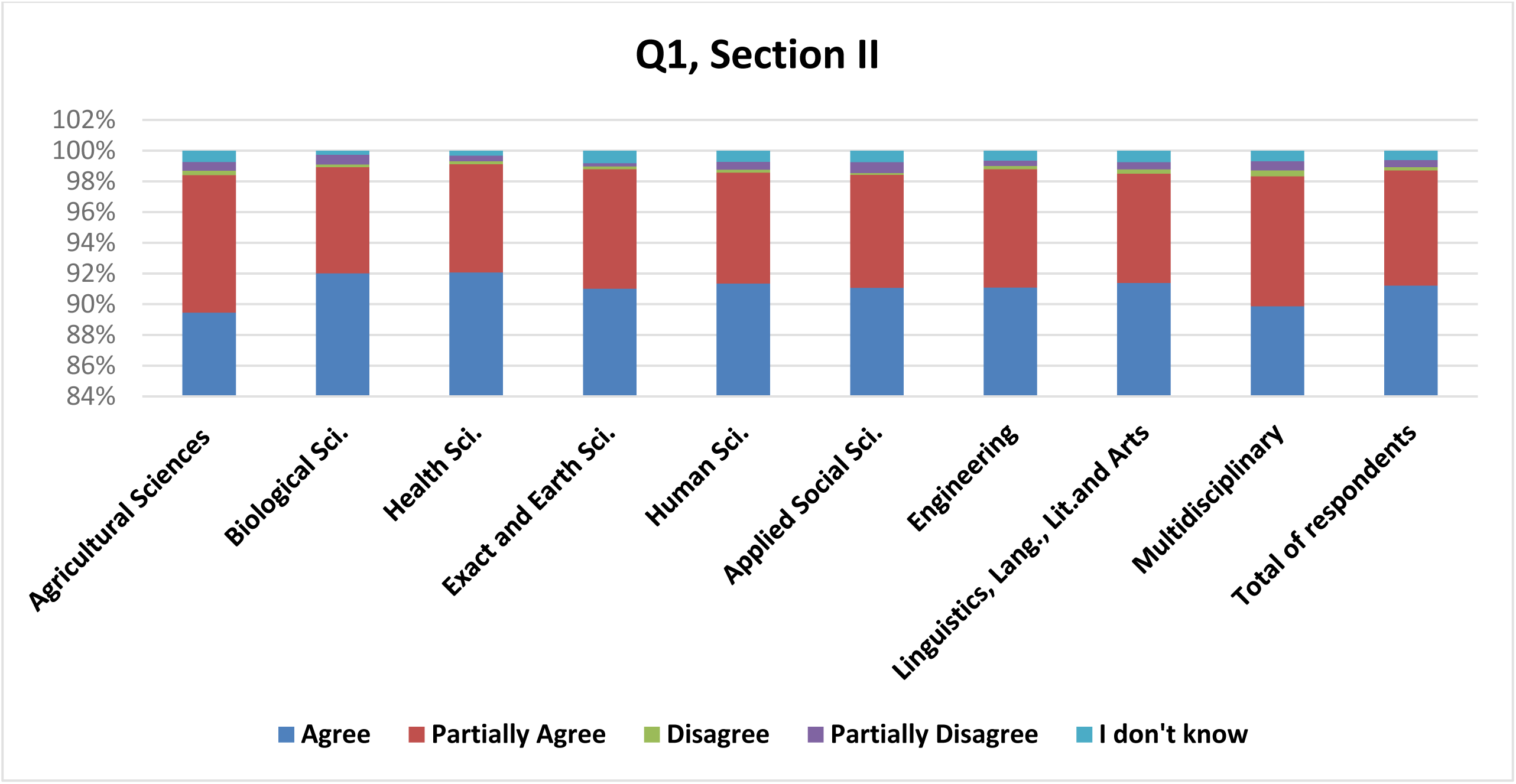
Patterns of response (n=25,105), according to field of knowledge, showing respondents’ level of agreement that plagiarism should be part of the research misconduct definition of US OSTP - **Q1, Section II** *The definition of misconduct established in 2000 by the U.S. Office of Science and Technology Policy (OSTP) is as follows: “Research misconduct is defined as fabrication, falsification, or plagiarism in proposing, performing, or reviewing research, or in reporting research results.” Thus, just as fabrication and falsification are research misconduct, so is plagiarism a form of research misconduct. Do you agree?*

**Fig. S2.**
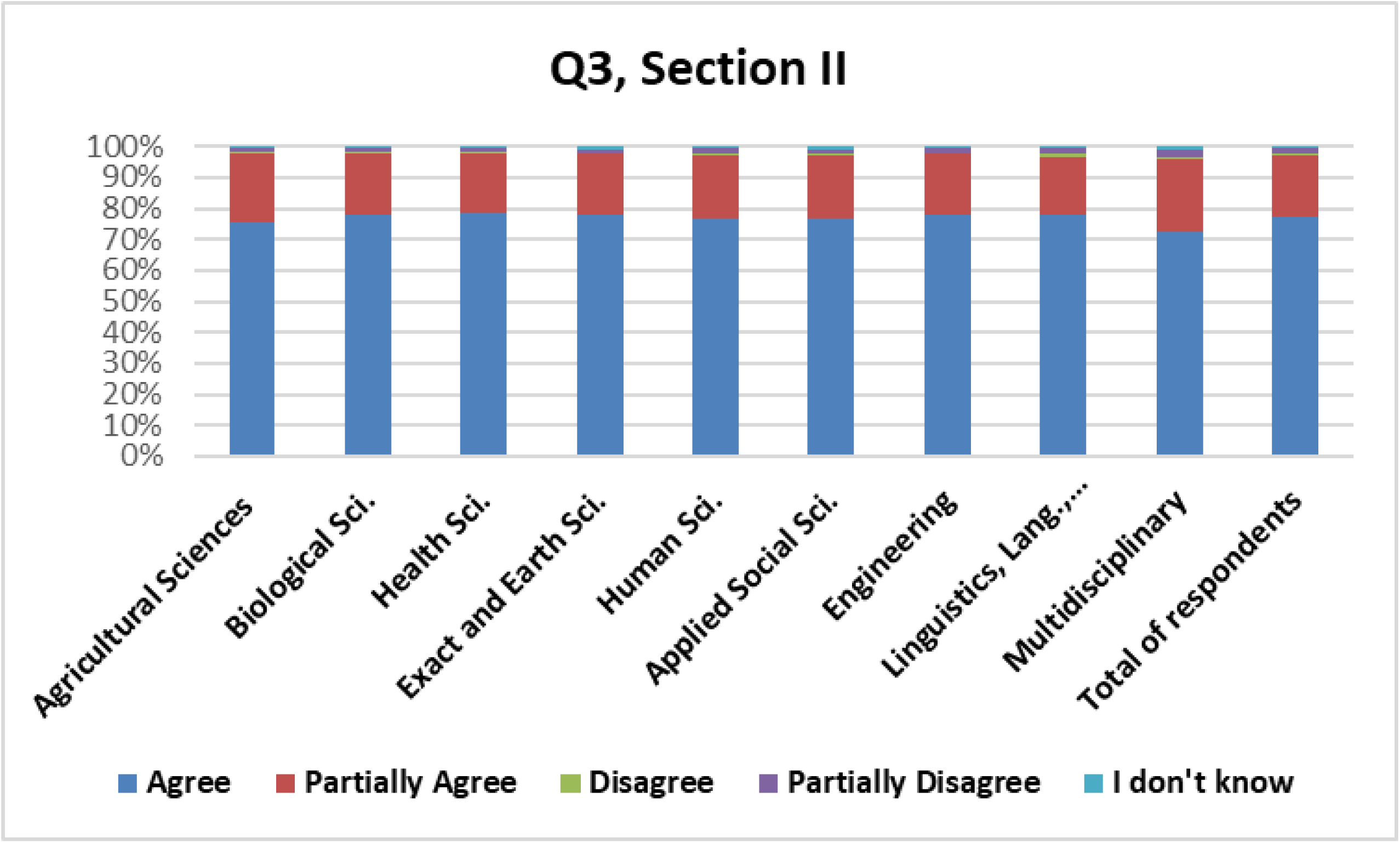
Patterns of response (n=24,993) in all fields for respondents’ level of agreement with the US OSTP’s definition of plagiarism - *The definition of plagiarism by OSTP, embraced by much of the international academic community, is as follows: “Appropriation of another person’s ideas, processes, results, or words without giving appropriate credit.”* **Q3, Section II** *Do you agree with this definition?*.

**Fig. S3.**
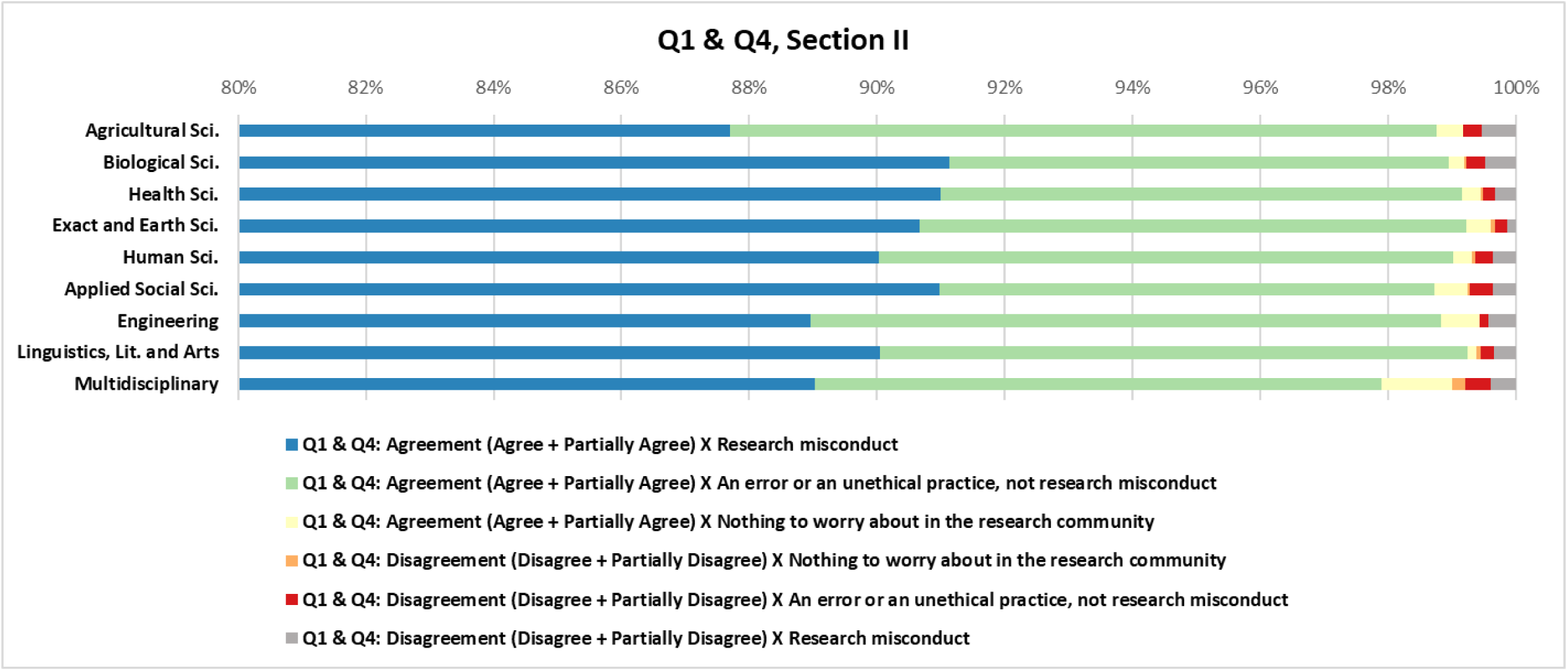
Patterns of response in all fields for respondents’ level of agreement (Agree + Partially Agree; Disagree + Partially Disagree) with **Q1, Section II,** US OSTP’s definition of plagiarism **& Q4, Section II –** Patterns of response across fields for *How do you see plagiarism in science? Nothing to worry about in the scientific community; An error, but not research misconduct; An unethical practice, but not research misconduct; Research misconduct; Research misconduct, except for textual plagiarism; Research misconduct, except for plagiarism of ideas*. The blue bar shows the percentages of agreement of respondents in each field with **Q1, Section II and** Research misconduct in **Q4, Section II**.

**Fig. S4.**
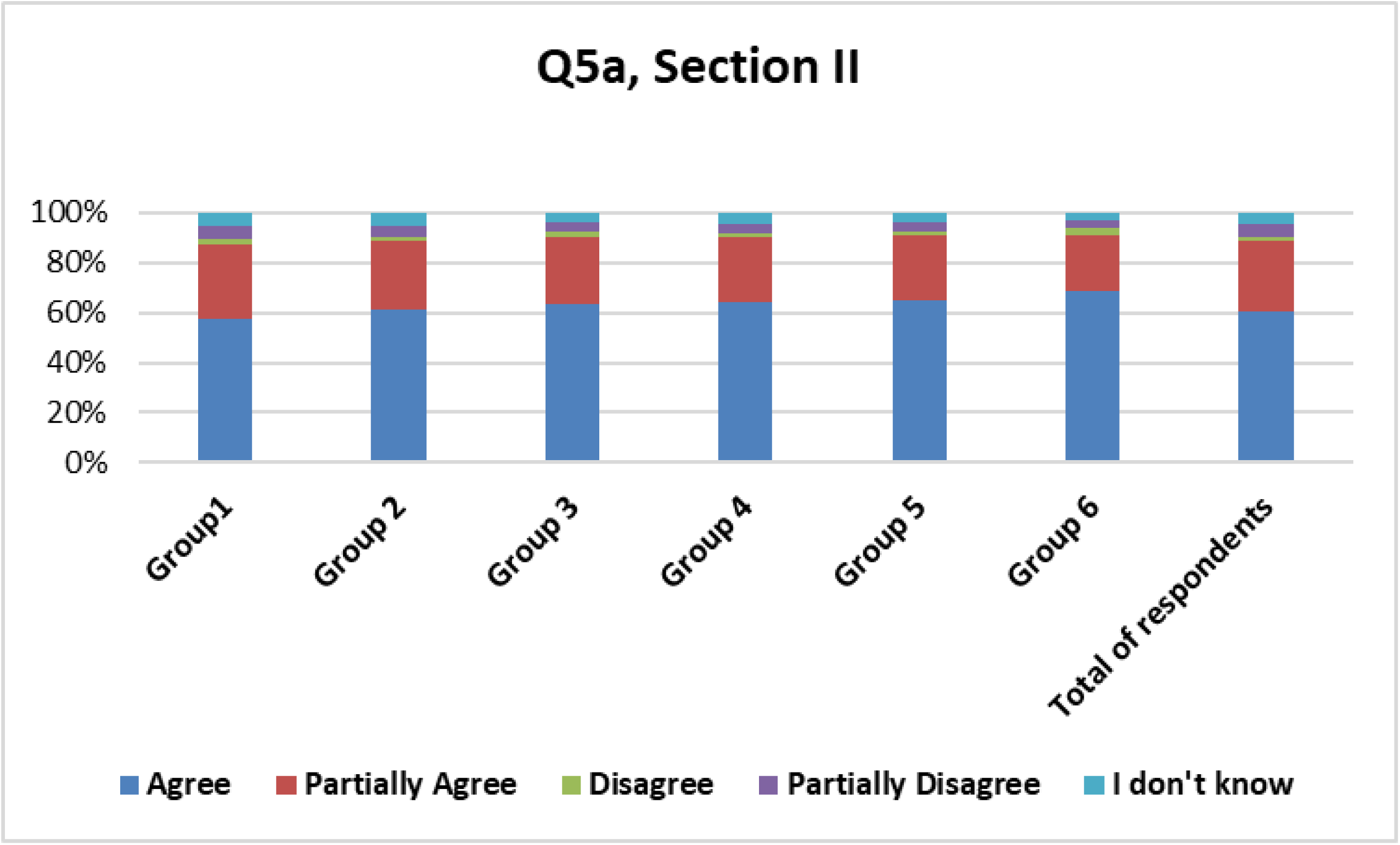
Patterns of response (n=25,029) according to time since earning the PhD: **Group1**: up to 5 years; **Group 2**: 5-10 years; **Group 3**: 10-15 years; **Group 4**: 15-20 years; **Group 5**: 20-25 years; **Group 6**: more than 25 years. Plagiarism and retractions - **Q5, Section II** *Recent surveys indicate an increase in plagiarism in scientific publications. Many of the cases have led to “retractions” of scientific papers (cancellation of publications). In this context, in 2010, Nature Reviews Genetics (NRG) retracted a review article for textual plagiarism [Nature Reviews Genetics 11:308(2010]. The plagiarism involved a single paragraph that had been paraphrased from an article submitted to Plant Science. The author of the NRG review was a referee for the Plant Science paper but she failed to cite it when she wrote the NRG review. In the retraction notice the NRG editors stated that the misappropriated paragraph was plagiarized and that the author of the NRG review had presented the ideas and hypotheses found in the original paragraph as if they were her own.* **Q5a, Section II** *Do you agree that textual plagiarism justifies a retraction*?.

**Fig. S5.**
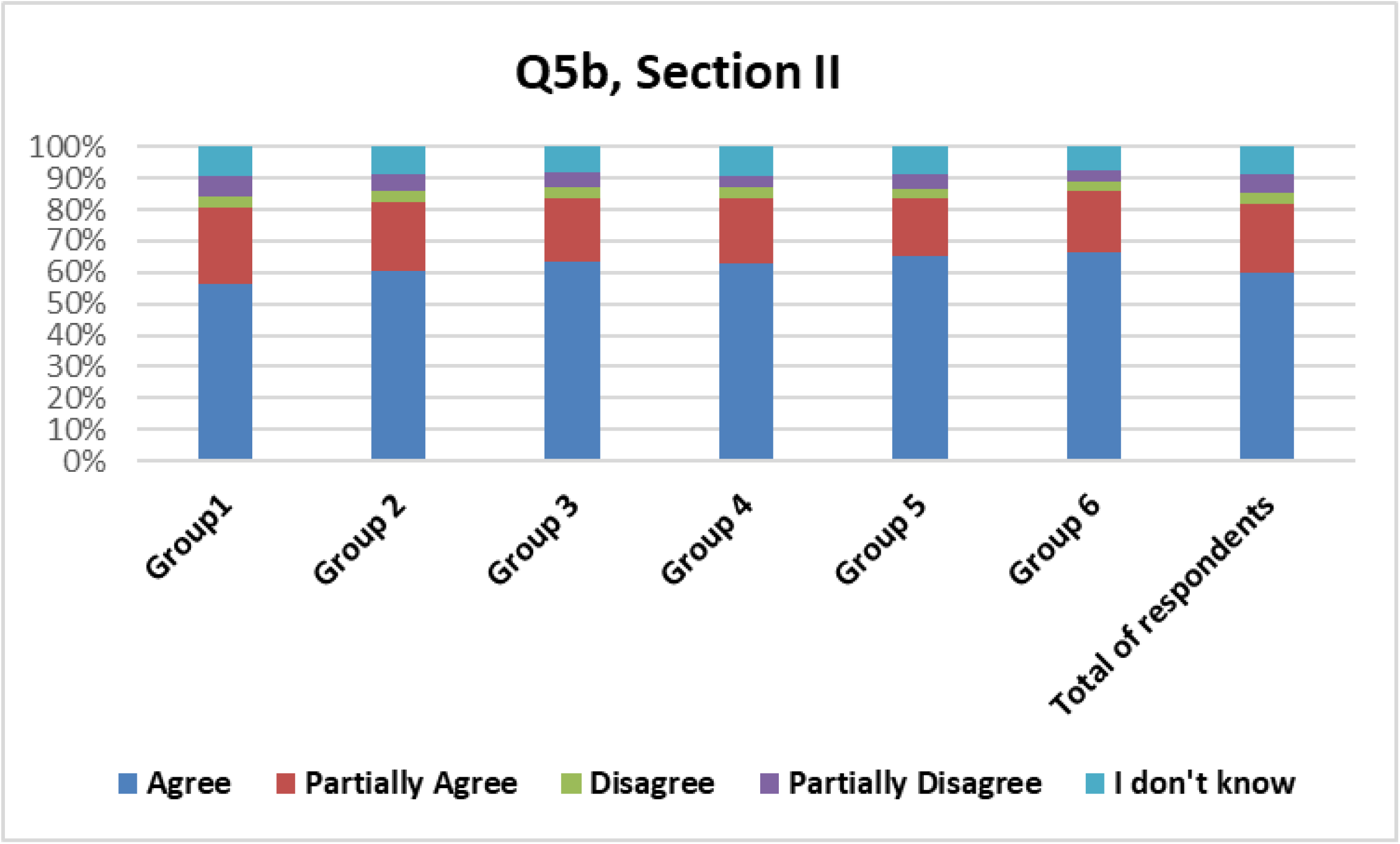
Patterns of response (n=24,996) according to time since earning the PhD: **Group1**: up to 5 years; **Group 2**: 5-10 years; **Group 3**: 10-15 years; **Group 4**: 15-20 years; **Group 5**: 20-25 years; **Group 6**: more than 25 years. Plagiarism and retractions (**Q5, Section II**). *Recent surveys indicate an increase in plagiarism in scientific publications. Many of the cases have led to “retractions” of scientific papers (cancellation of publications). In this context, in 2010, Nature Reviews Genetics (NRG) retracted a review article for textual plagiarism [Nature Reviews Genetics 11:308(2010]. The plagiarism involved a single paragraph that had been paraphrased from an article submitted to Plant Science. The author of the NRG review was a referee for the Plant Science paper but she failed to cite it when she wrote the NRG review. In the retraction notice the NRG editors stated that the misappropriated paragraph was plagiarized and that the author of the NRG review had presented the ideas and hypotheses found in the original paragraph as if they were her own*. **Q5b**, **Section II** *Do you agree with the retraction in this case?*.

**Fig. S6.**
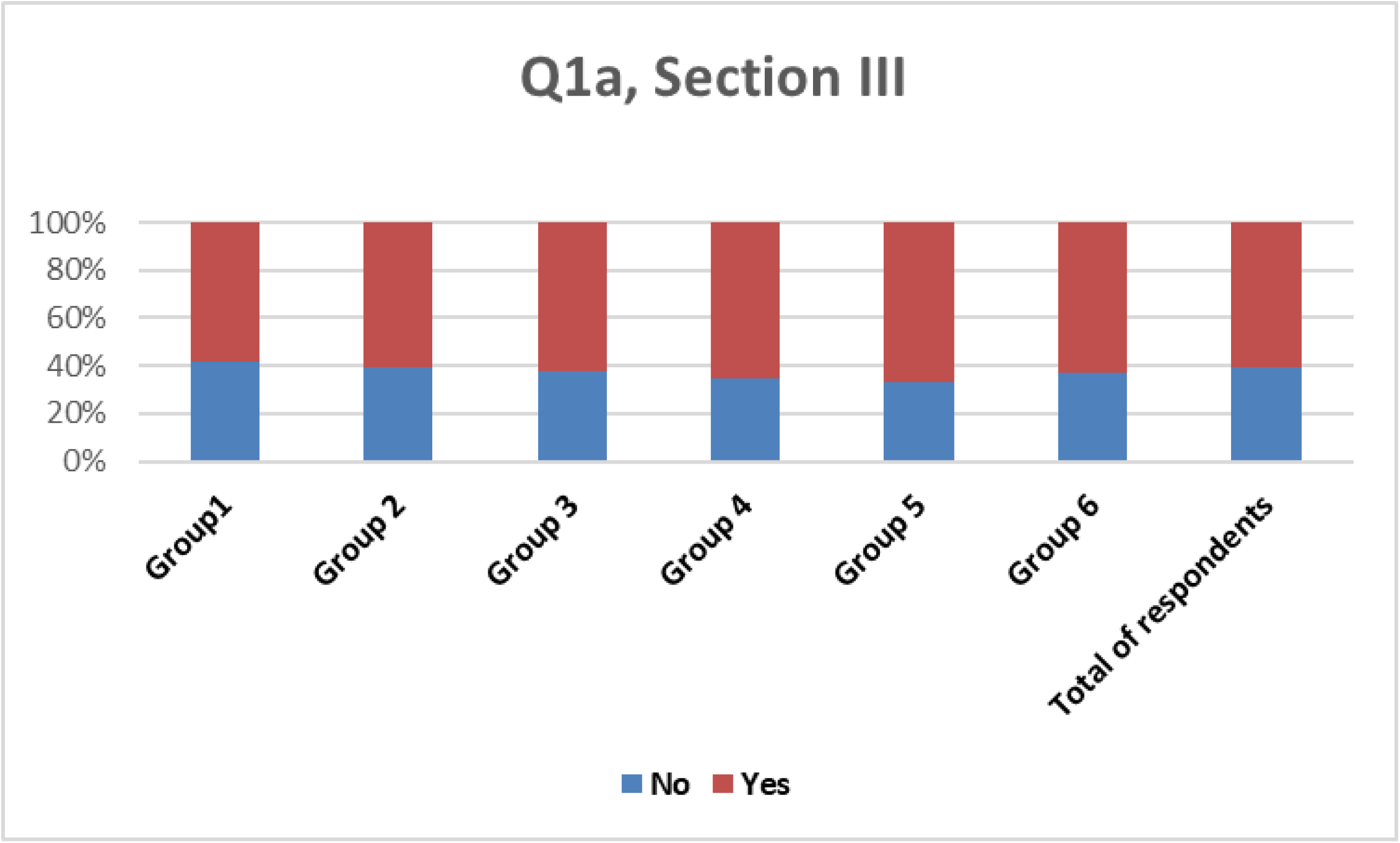
Patterns of response (n=24,430) across all fields for respondents’ awareness of plagiarism detection software used in the publication system. **Q1a, Section III** *Have you ever heard of such software used by scientific publishers?*.

**Fig. S7.**
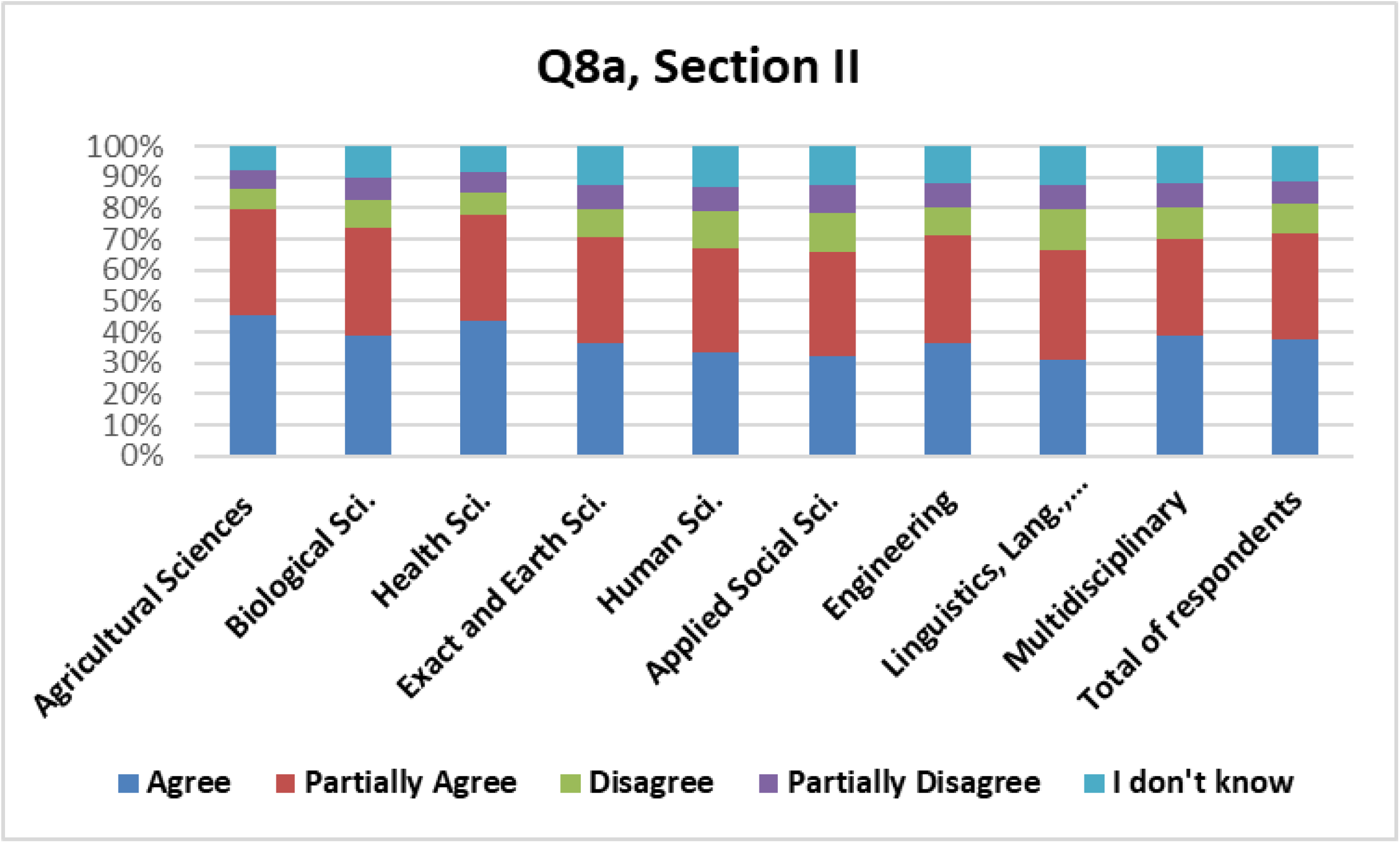
Patterns of response (n= 25,008) across fields for **Q8, Section II** *In the context of plagiarism, it is worth noting that in Brazil:* **Q8a, Section II** *Graduate students are not familiar with the international concept of academic plagiarism*.

**Fig. S8.**
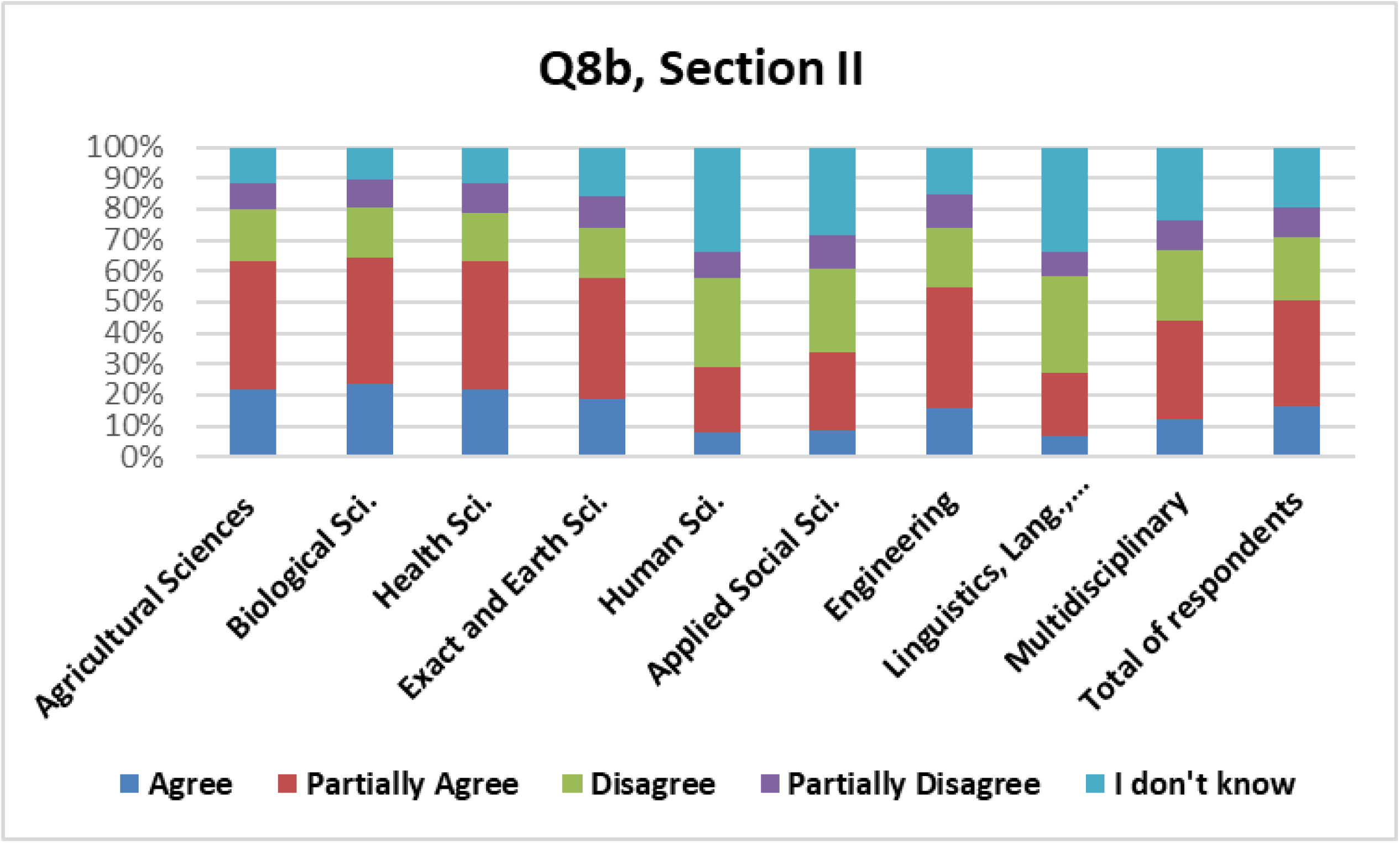
Patterns of response (n=25,021) across fields for **Q8, Section II -** *In the context of plagiarism, it should be highlighted that for Brazil* - **Q8b, Section II** *Graduate students tend to engage in textual plagiarism in scientific articles in English because they are not fluent in English*.

**Table S1.**
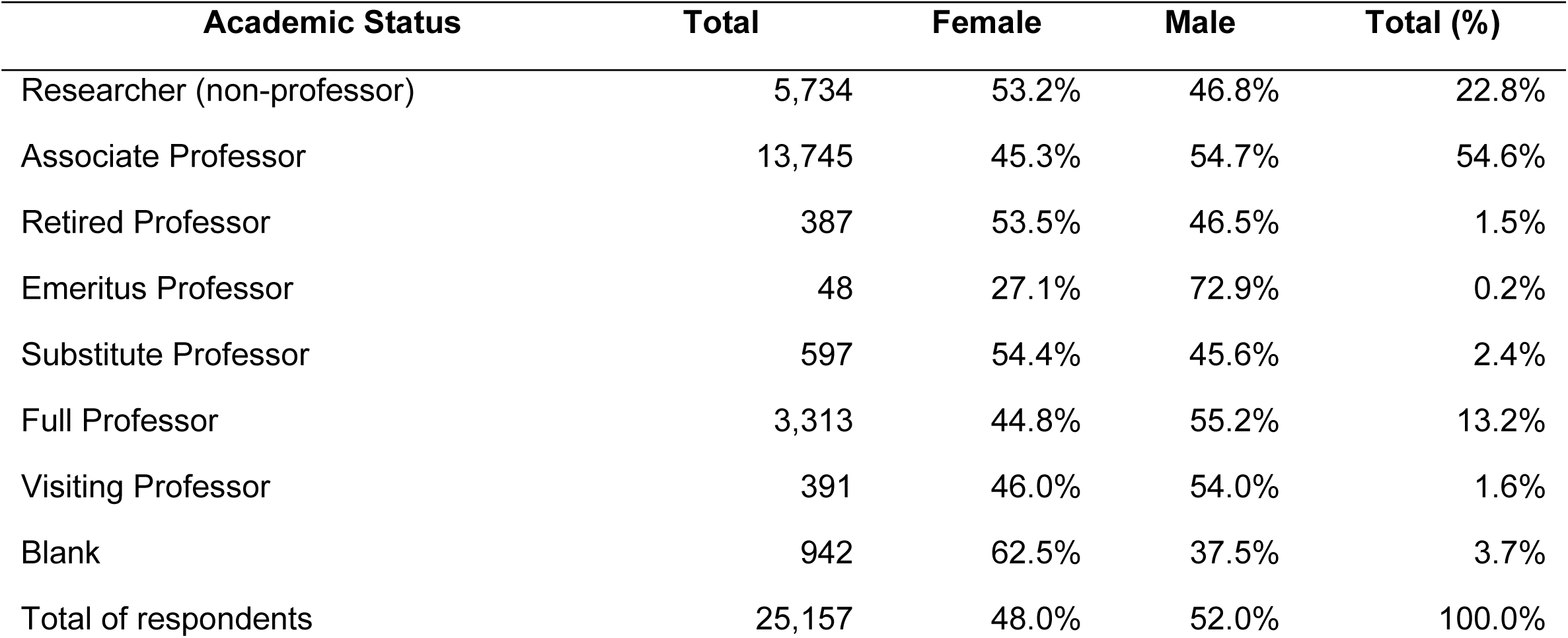
PhD holders responding to the national survey (n=25,157) by academic status in 2013 - self-declared information in 2014

**Table S2.** Plagiarism and retractions - **Q5, Section II** *Recent surveys indicate an increase in plagiarism in scientific publications. Many of the cases have led to “retractions” of scientific papers (cancellation of publications). In this context, in 2010, Nature Reviews Genetics (NRG) retracted a review article for textual plagiarism [Nature Reviews Genetics 11:308(2010]. The plagiarism involved a single paragraph that had been paraphrased from an article submitted to Plant Science. The author of the NRG review was a referee for the Plant Science paper but she failed to cite it when she wrote the NRG review. In the retraction notice the NRG editors stated that the misappropriated paragraph was plagiarized and that the author of the NRG review had presented the ideas and hypotheses found in the original paragraph as if they were her own*. **Q5b, Section II** *Do you agree with retraction in this particular case*? (n=24,996)

| Field of knowledge | Agree | Partially Agree | Disagree | Partially Disagree | I don't know |
| --- | --- | --- | --- | --- | --- |
| Agricultural Sciences | 58.7% | 24.7% | 3.8% | 6.1% | 6.7% |
| Biological Sciences | 58.2% | 24.2% | 4.5% | 6.2% | 6.9% |
| Health Sciences | 60.0% | 23.7% | 3.5% | 6.8% | 6.0% |
| Exact and Earth Sciences | 56.0% | 22.8% | 4.3% | 5.9% | 11.0% |
| Human Sciences | 63.6% | 19.4% | 2.7% | 4.8% | 9.5% |
| Applied Social Sciences | 62.9% | 20.2% | 3.0% | 4.6% | 9.4% |
| Engineering | 55.7% | 23.3% | 3.8% | 5.7% | 11.5% |
| Linguistics, Language & Literature and Arts | 62.6% | 21.0% | 2.7% | 4.5% | 9.2% |
| Multidisciplinary | 59.4% | 21.9% | 3.5% | 5.3% | 9.9% |
| Total of respondents | 59.6% | 22.4% | 3.6% | 5.7% | 8.7% |

**Table S3.** Plagiarism and graduate students – **Q7, Section II** *Have you ever encountered any case of plagiarism (partial or total) by any graduate student (not necessarily from your Program) in the last four years*? – **Q7a** - *When reviewing a Master’s thesis* (n=24,830)

| Field of knowledge | No | Yes,<br>one<br>case | Yes,<br>two<br>cases | Yes,<br>three<br>cases | Yes,<br>more<br>than<br>three<br>cases | Yes,<br>many<br>cases |
| --- | --- | --- | --- | --- | --- | --- |
| Agricultural Sciences | 50.5% | 20.9% | 9.4% | 2.1% | 5.3% | 11.9% |
| Biological Sciences | 59.8% | 19.8% | 7.7% | 1.4% | 4.3% | 7.2% |
| Health Sciences | 57.9% | 20.6% | 8.0% | 1.3% | 4.0% | 8.1% |
| Exact and Earth Sciences | 65.2% | 16.7% | 6.1% | 1.4% | 3.5% | 7.1% |
| Health Sciences | 50.4% | 21.9% | 9.9% | 2.7% | 4.4% | 10.6% |
| Applied Social Sciences | 50.1% | 20.9% | 9.5% | 2.7% | 5.1% | 11.8% |
| Engineering | 61.3% | 18.3% | 6.9% | 1.9% | 3.9% | 7.7% |
| Linguistics, Language & Literature and Arts | 53.2% | 20.6% | 8.6% | 2.6% | 5.0% | 10.0% |
| Multidisciplinary | 51.6% | 18.7% | 9.5% | 2.7% | 5.3% | 12.1% |
| Total of respondents | 56.3% | 19.9% | 8.2% | 1.9% | 4.4% | 9.2% |

**Table S4.** Plagiarism and graduate students – **Q7, Section II** *Have you ever encountered any case of plagiarism (partial or total) of any graduate student (not necessarily from your Program) in the last four years?* – **Q7b** *When reviewing a PhD thesis* (n=24,329)

| Field of knowledge | No | Yes,<br>one<br>case | Yes,<br>two<br>cases | Yes,<br>three<br>cases | Yes,<br>more<br>than<br>three<br>cases | Yes,<br>many<br>cases |
| --- | --- | --- | --- | --- | --- | --- |
| **Agricultural Sciences** | **66.0%** | **16.6%** | **5.4%** | **1.1%** | **3.5%** | **7.4%** |
| **Biological Sciences** | **75.6%** | **13.5%** | **3.6%** | **0.8%** | **2.4%** | **4.1%** |
| **Health Sciences** | **79.3%** | **11.7%** | **3.1%** | **0.8%** | **1.8%** | **3.4%** |
| **Exact and Earth Sciences** | **80.8%** | **10.2%** | **3.0%** | **0.8%** | **1.6%** | **3.6%** |
| **Health Sciences** | **78.2%** | **12.9%** | **3.2%** | **0.8%** | **1.4%** | **3.4%** |
| **Applied Social Sciences** | **81.2%** | **10.7%** | **2.8%** | **0.9%** | **2.0%** | **2.4%** |
| **Engineering** | **79.3%** | **11.9%** | **2.8%** | **0.7%** | **1.9%** | **3.4%** |
| **Linguistics, Language & Literature and Arts** | **77.3%** | **12.9%** | **3.0%** | **0.7%** | **2.0%** | **4.1%** |
| **Multidisciplinary** | **76.7%** | **12.8%** | **3.3%** | **0.5%** | **2.1%** | **4.6%** |
| **Total of respondents** | **77.5%** | **12.4%** | **3.3%** | **0.8%** | **2.0%** | **3.9%** |

We asked a series of questions exploring views of originality in the writing of a research article (**Fig.S9, S10, S11, S12**). Each question poses a specific situation related to attribution of credit in the process of borrowing from the literature. The series of questions is the following:

Plagiarism and originality in the writing of a research article should be questioned if the author used entire paragraphs from other works without a citation (**Q4a, Section III);** Plagiarism and originality in the writing of a research article should be questioned if the author used entire paragraphs from other works with a citation but without quotation marks (**Q4b, Section III**); Plagiarism and originality in the writing of a research article should be questioned if the author used paragraphs from other works correctly paraphrased, but without citation. (**Q4c, Section III**); Plagiarism and originality in the writing of a research article should be questioned if the author used entire paragraphs from other works without citation, considering that he/she was a co-author in this previous publication. (**Q4d, Section III**).

**Fig. S9.**
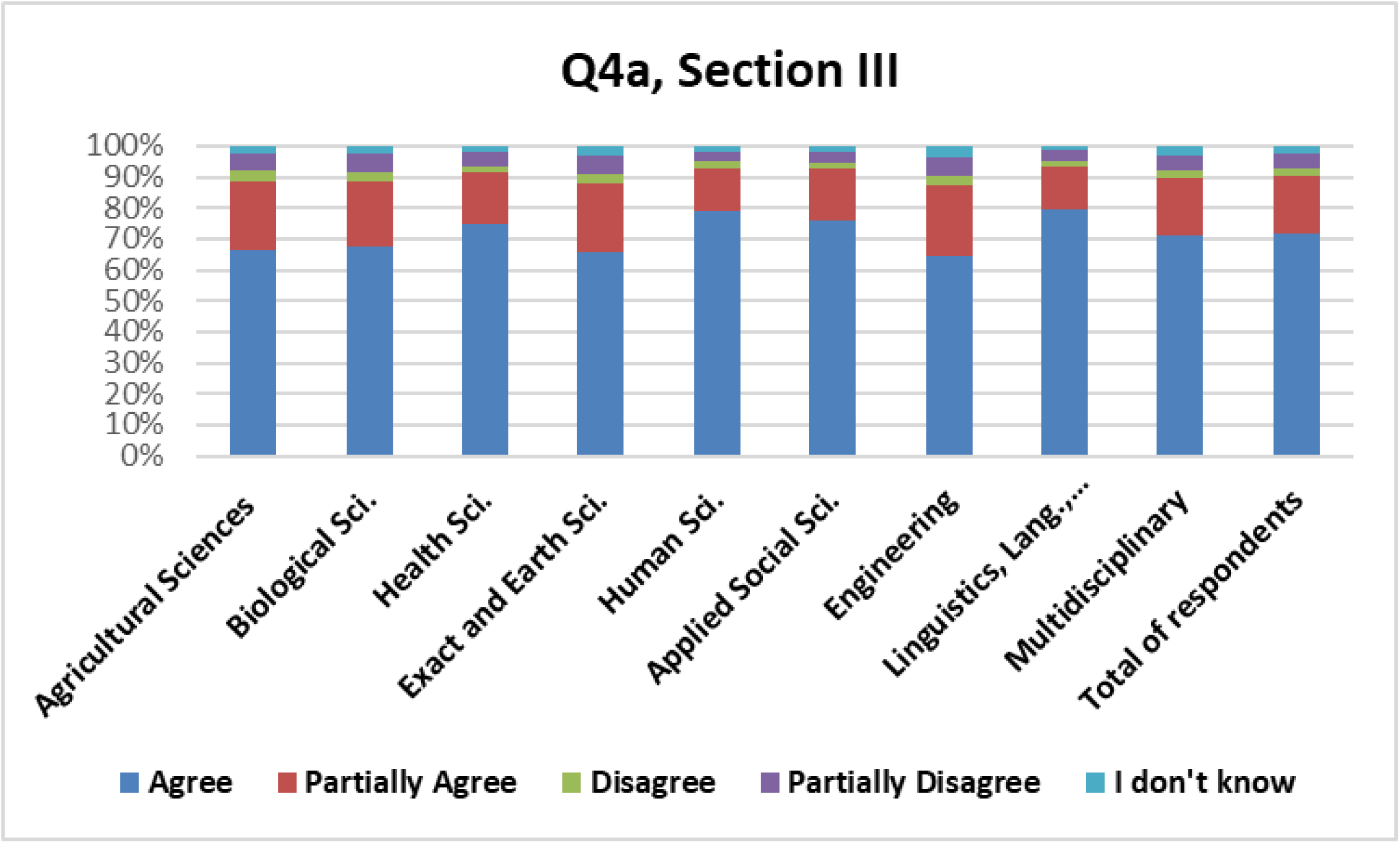
Patterns of response (n=24,323) for **Q4a, Section III** *The originality of the results in a research paper should be questioned if the author of that paper **copied entire paragraphs without citation from others’ previously published papers.***

**Fig. S10.**
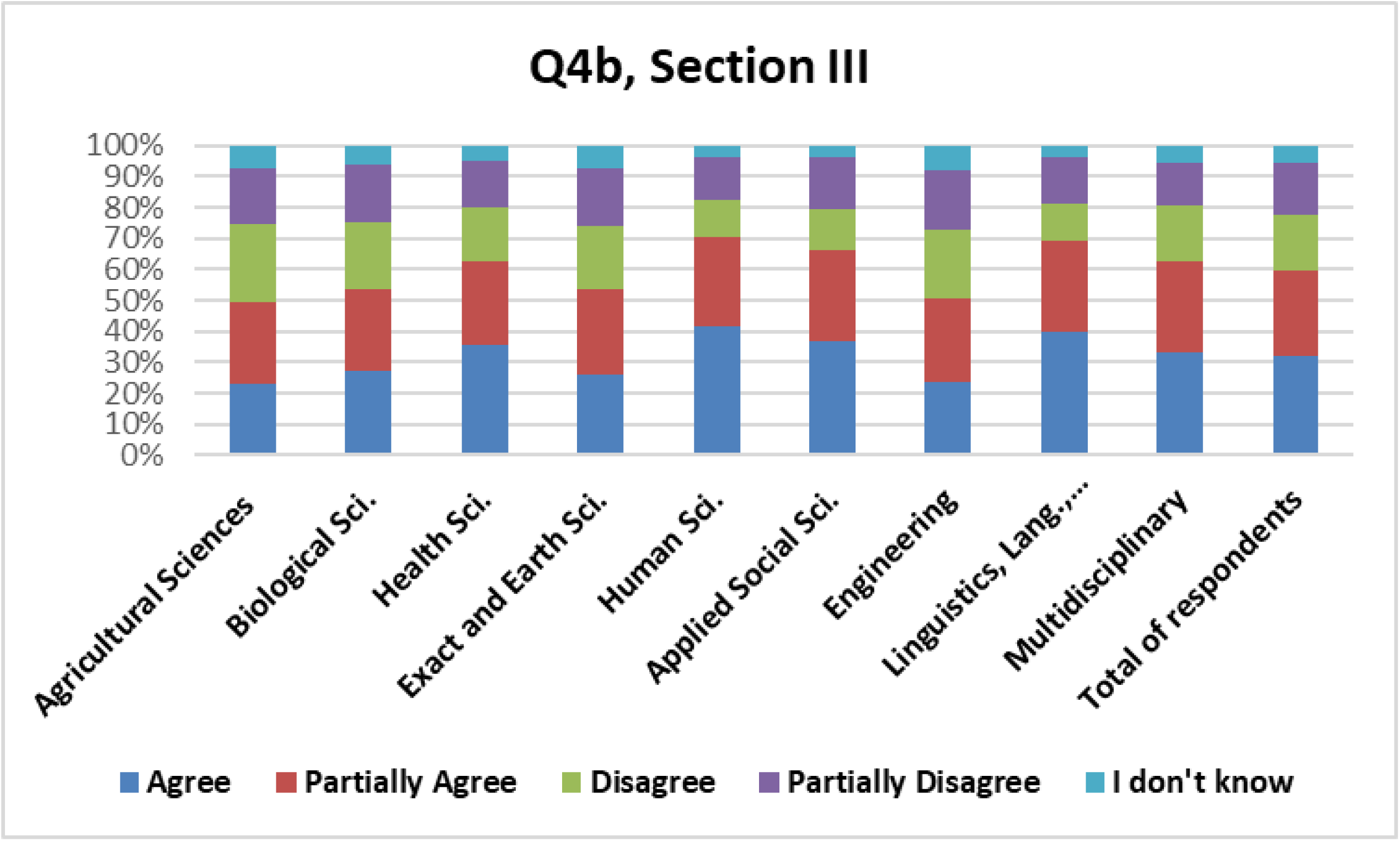
Patterns of response (n=24,336) for **Q4b, Section III** *The originality of the results in a research paper should be questioned if the author of that paper **copied entire paragraphs from others’ previously published papers, citing these sources but without enclosing the copied text in quotation marks.***

**Fig. S11.**
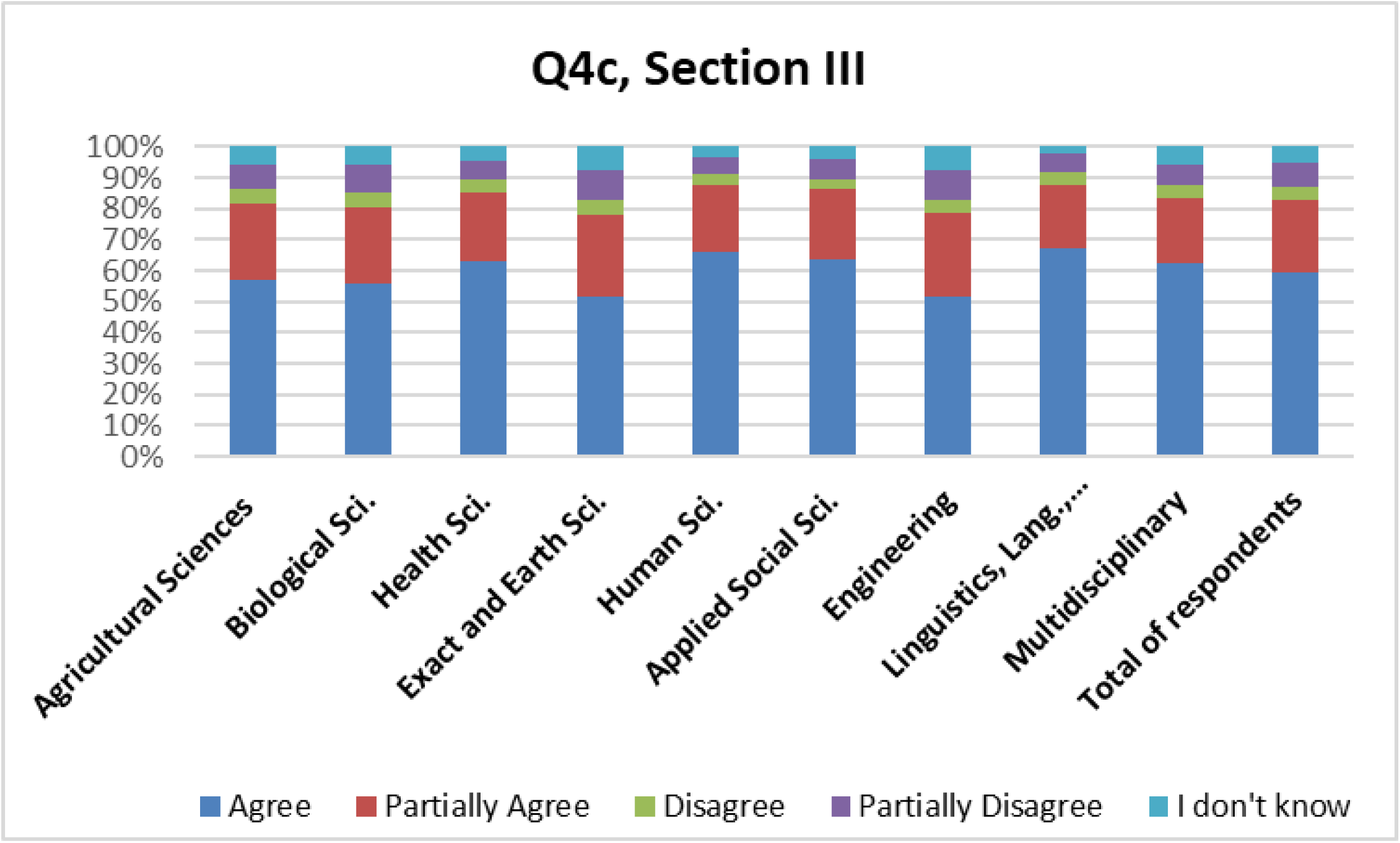
Patterns of response (n=24,259) for **Q4c, Section III** *The originality of the results in a research paper should be questioned if the author of that paper correctly paraphrased **entire paragraphs from others’ previously published papers but without citing the original sources.***

**Fig. S12.**
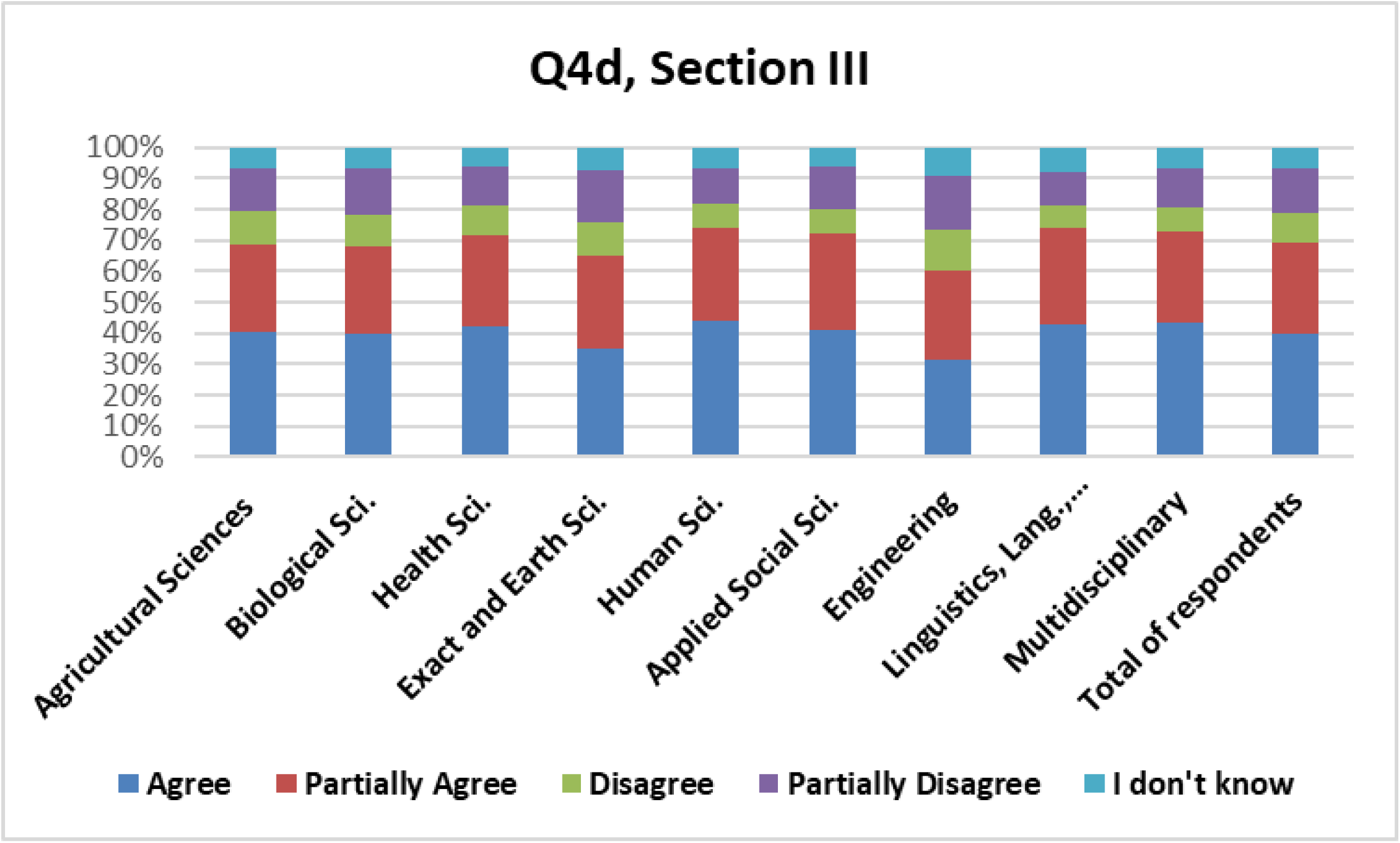
Patterns of response (n=24,392) for **Q4d, Section III** *The originality of the results in a research paper should be questioned if the author of that paper copied **entire paragraphs without citation from other previously published papers, and that same person was a co-author of all the publications involved.***

Note that whereas widespread agreement is noted for **Q4a**, views are stricter for Human Sciences, Applied Social Sciences, Linguistics, Language & Literature and Arts, and Health Sciences. Similar response patterns for these fields are also found for **Q4b**, including the Multidisciplinary group (*SI 2*). Human Sciences, Applied Social Sciences, Linguistics, Language & Literature and Arts, and Health Sciences are the fields with the smallest percentages for “I don’t know” (4%), while the highest percentage is for Engineering (8%). Similar patterns also emerge for **Q4c.** Nevertheless, as corroborated by patterns shown for **Q4a, Section III**, differences are less noticeable for questions addressing formal definitions of research misconduct and plagiarism, such as **Q1**, **Q2a and Q2b,** and **Q3**, **Section II**. Overall, particularly for questions that imply the application of the concept of plagiarism to real-case scenarios, the Human Sciences, Applied Social Sciences, Linguistic & Literature and Arts, as well as the Health Sciences show similar response patterns (*SI 2*).

Kruskal-Wallis non-parametric statistical test (**1**) was used to verify the null hypothesis – distribution of response patterns would be the same across the nine *grand* fields.

Considering that there was no significant difference among response patterns for these *grand* fields, p-values were calculated for questions in **Section II and Section III**. For p-values smaller than 0.05, the difference was considered statistically significant, and the null hypothesis was rejected. We then carried out post-hoc tests for pairwise comparisons (Sample 1-Sample 2) to see which pair would differ significantly. We then list the adjusted p-values for each. In each diagram, the orange line joining specific groups indicates statistically significant differences.

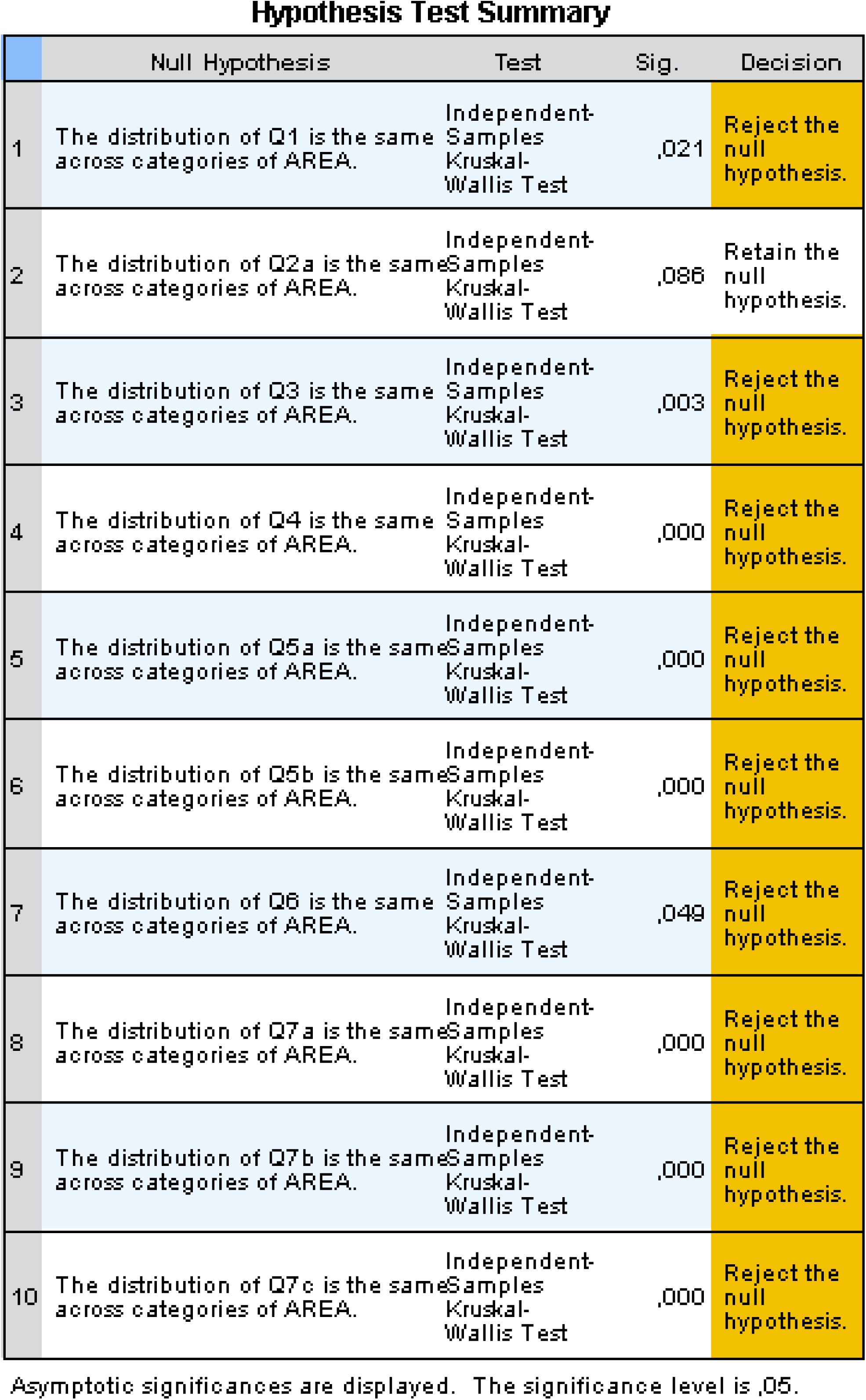
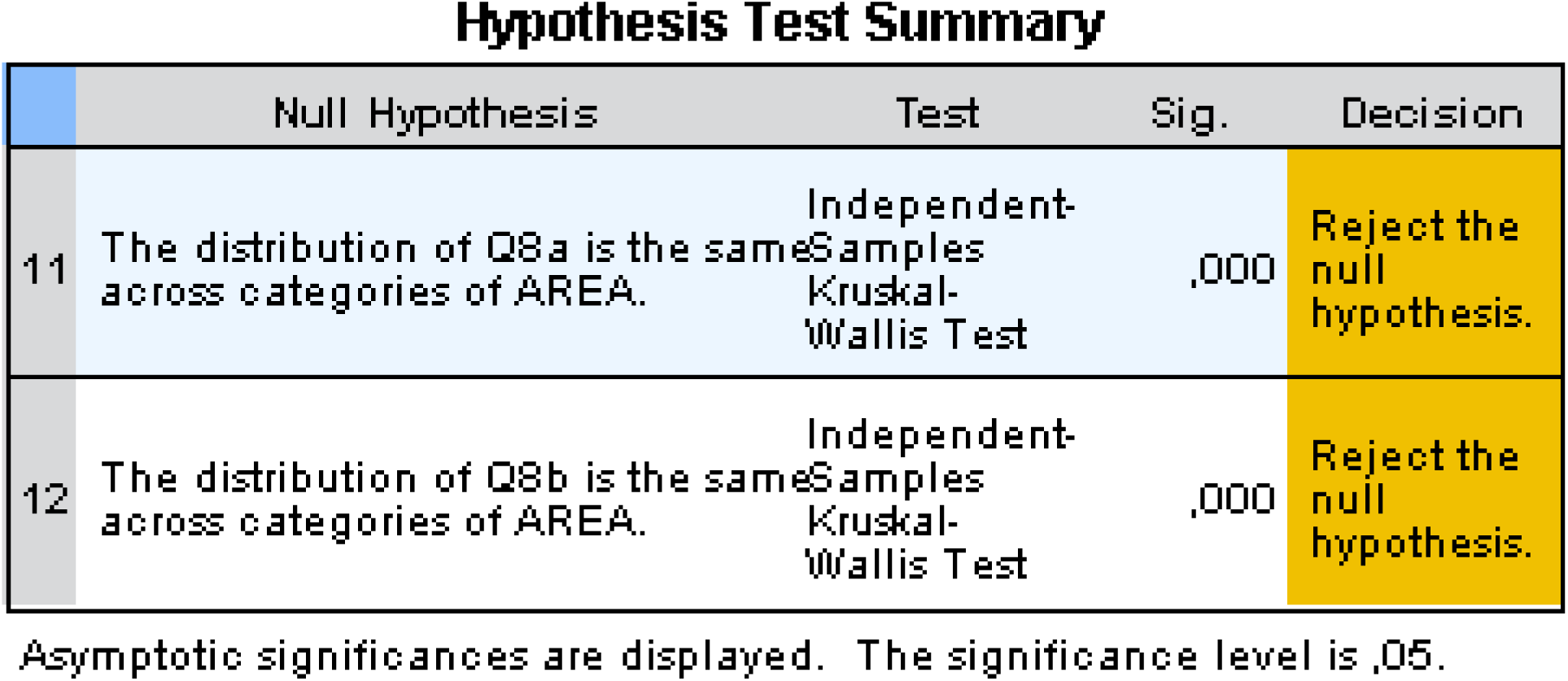

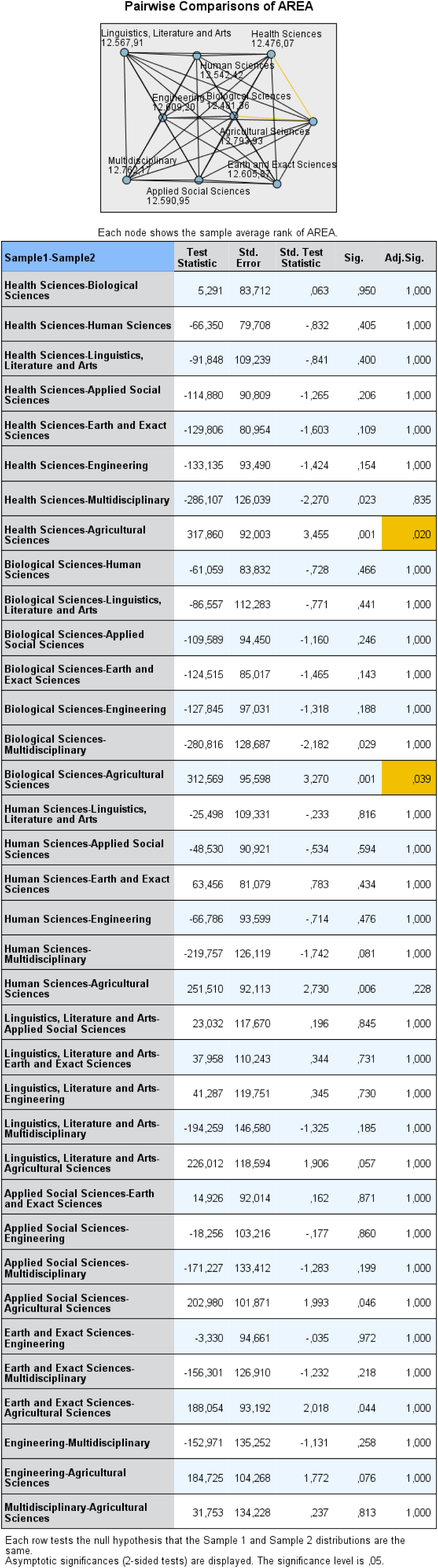

From the adjusted significance level, for Q1, **Section II**, significant differences in response patterns were found for the following fields:

- **Health Sciences and** Agricultural Sciences.
- **Biological Sciences and** Agricultural Sciences.

For Q2a, no significant differences were found.

For Q3, see below:

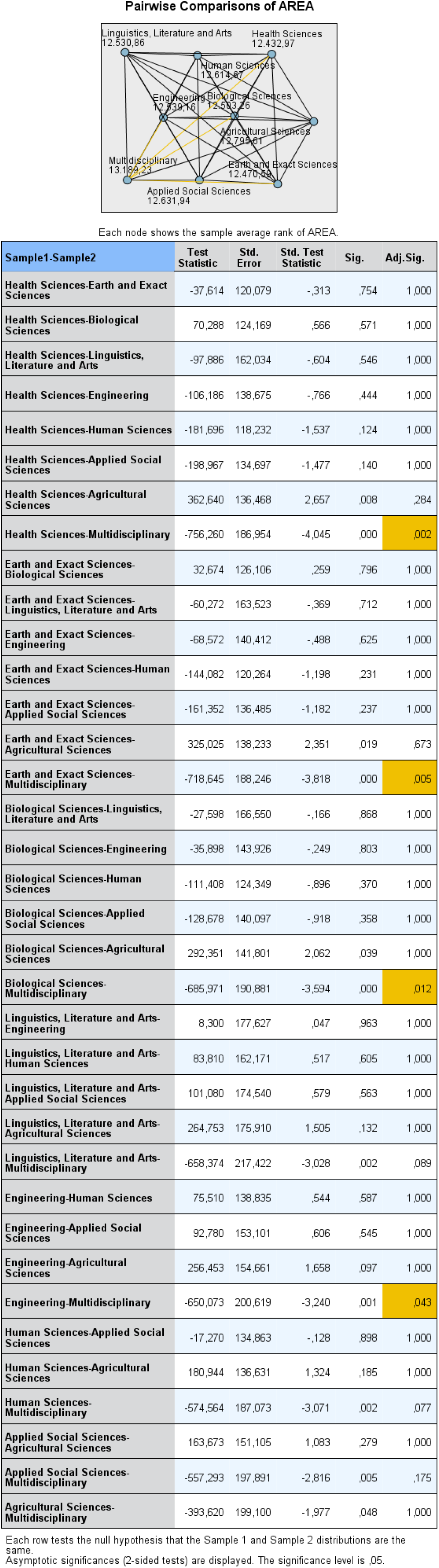

For Q3, significant differences in response patterns were observed between the following fields:

- **Health Sciences and** Multidisciplinary.
- **Earth and Exact Sciences and** Multidisciplinary.
- **Biological Sciences and** Multidisciplinary.
- **Engineering and** Multidisciplinary.

For Q4, see below:

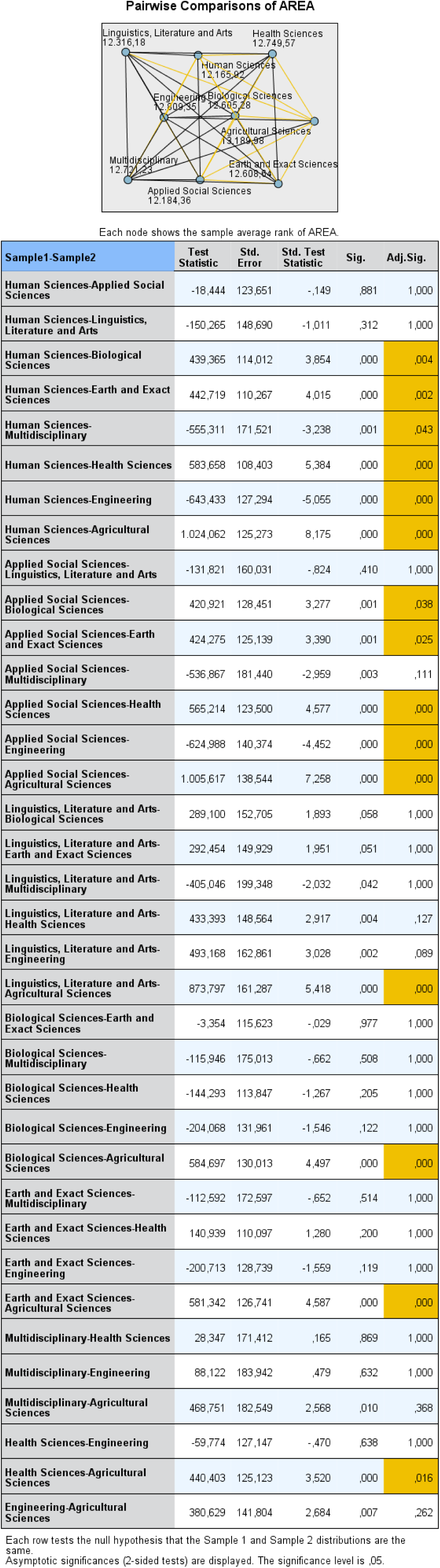

For Q4, significant differences in response patterns were observed between the following fields:

- **Human Sciences and** Biological Sciences; and Earth and Exact Sciences; and Multidisciplinary; and Health Sciences; and Engineering; and Agricultural Sciences.
- **Applied Social Sciences and** Biological Sciences; and Earth and Exact Sciences; and Health Sciences; and Engineering; and Agricultural Sciences.
- **Linguistics, Literature and Arts and** Agricultural Sciences.
- **Biological Sciences and** Agricultural Sciences.
- **Earth and Exact Sciences and** Agricultural Sciences.
- **Health Sciences and** Agricultural Sciences.

For Q5a, see below:

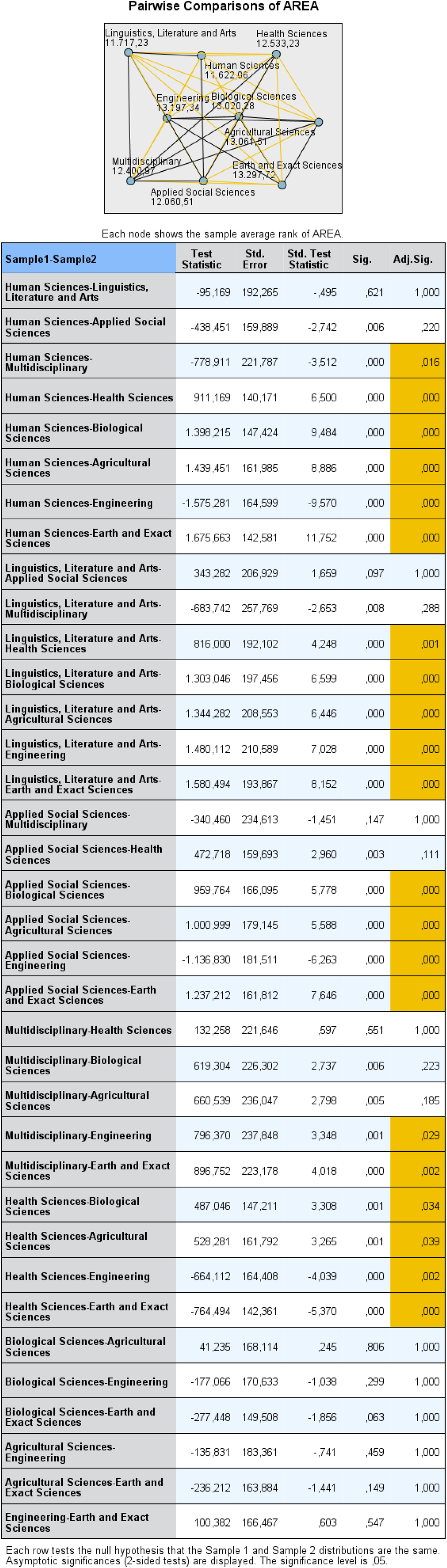

For Q5a, significant differences in response patterns were observed between the following fields:

- **Human Sciences and** Multidisciplinary; and Health Sciences; and Biological Sciences; and Agricultural Sciences; and Engineering and Earth and Exact Sciences.
- **Linguistics, Literature and Arts and** Health Sciences; and Biological Sciences; and Agricultural Sciences; and Engineering; and Earth and Exact Sciences.
- **Applied Social Sciences and** Biological Sciences; and Agricultural Sciences; and Engineering; and Earth and Exact Sciences.
- **Multidisciplinary and** Engineering; and Earth and Exact Sciences;
- **Health Sciences and** Biological Sciences; and Agricultural Sciences; and Engineering; and Earth and Exact Sciences.

For Q5b, see below:

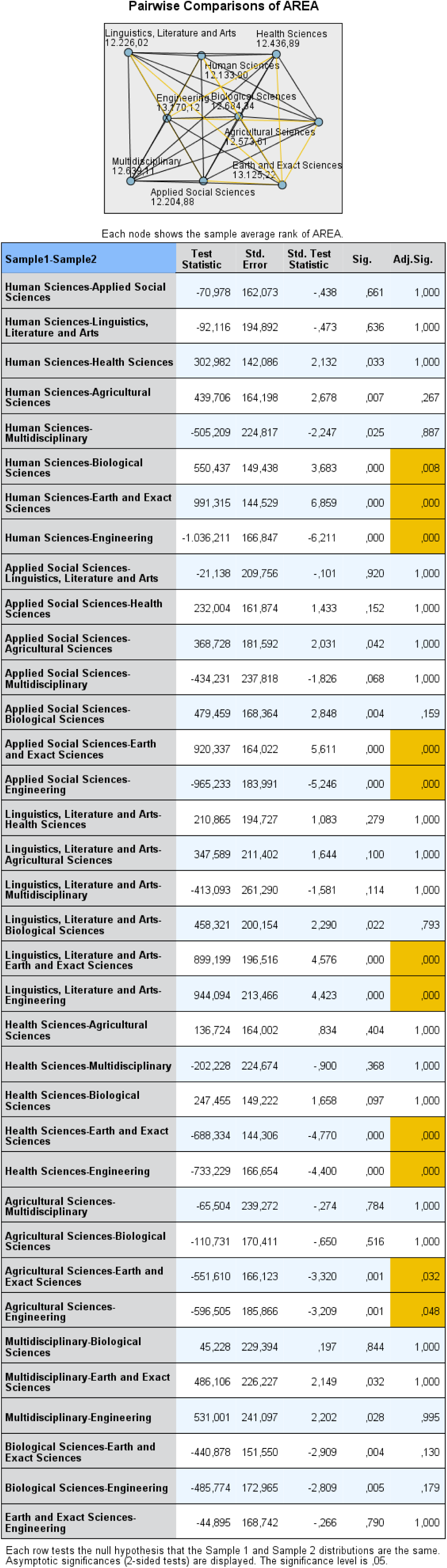

For Q5b, significant differences in response patterns were observed between the following fields:

- **Human Sciences and** Biological Sciences; and Earth and Exact Sciences; and Engineering.
- **Applied Social Sciences and** Earth and Exact Sciences; and Engineering.
- **Linguistics, Literature and Arts and**; Earth and Exact Sciences; and Engineering.
- **Health Sciences and**; Earth and Exact Sciences; and Engineering.
- **Agricultural Sciences and**; and Earth and Exact Sciences; and Engineering.

For Q6, see below:

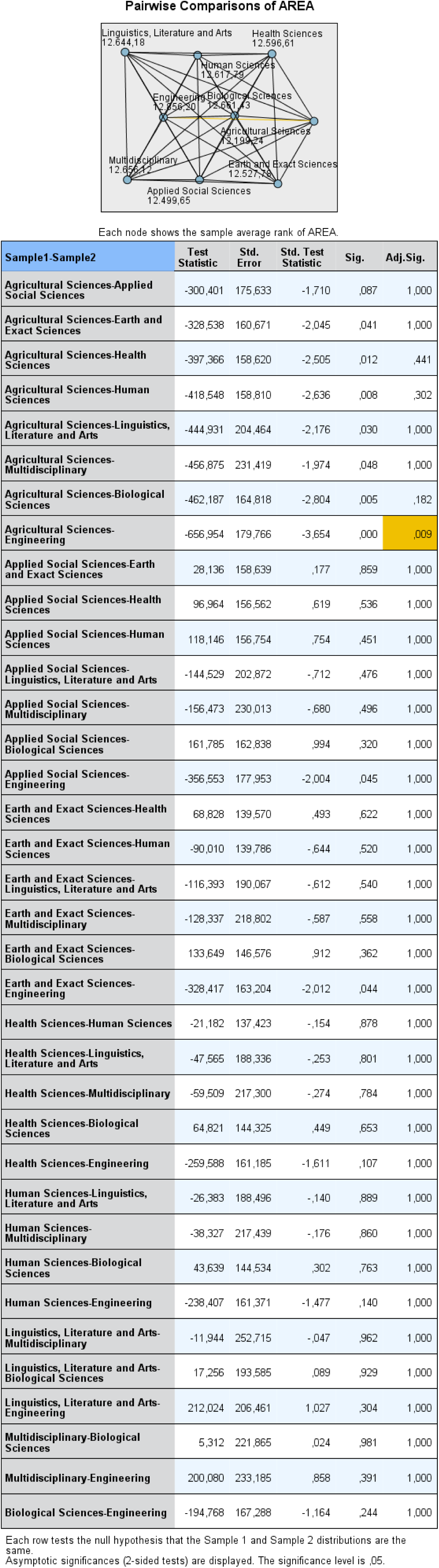

For Q6, the only significant differences in response patterns were observed between the following fields:

- **Agricultural Sciences and** Engineering.

– Note however, that this question is about being aware of a particular document and that would not involve views or perceptions.

For Q7a, see below:

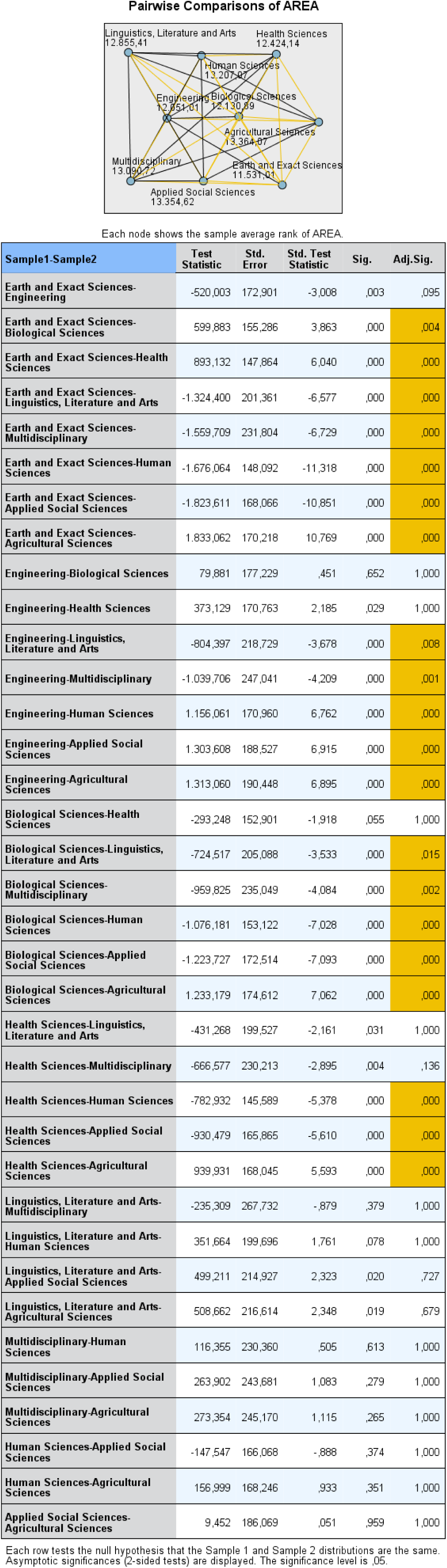

For Q7a, significant differences in response patterns were observed between the following fields:

- **Earth and Exact Sciences and** Biological Sciences; and Health Sciences; Linguistics, Literature and Arts; and Multidisciplinary; and Human Sciences; and Applied Social Sciences; and Agricultural Sciences.
- **Engineering and** Linguistics, Literature and Arts; and Multidisciplinary; and Human Sciences; and Applied Social Sciences; and Agricultural Sciences.
- **Biological Sciences and** Linguistics, Literature and Arts; and Multidisciplinary; and Human Sciences; and Applied Social Sciences; and Agricultural Sciences.
- **Health Sciences and** Human Sciences; and Applied Social Sciences; and Agricultural Sciences.

For Q7b, see below:

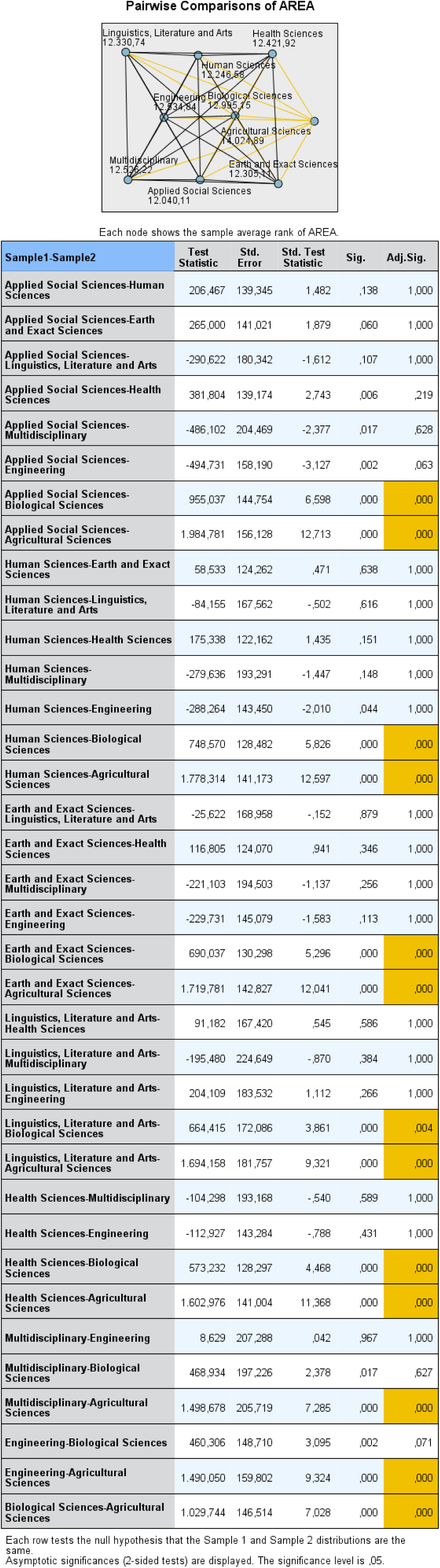

For Q7b, significant differences in response patterns were observed between the following fields:

- **Applied Social Sciences and** Biological Sciences; and Agricultural Sciences.
- **Human Sciences and** Biological Sciences; and Agricultural Sciences.
- **Earth and Exact Sciences and** Biological Sciences; and Agricultural Sciences.
- **Linguistics, Literature and Arts and** Biological Sciences; and Agricultural Sciences.
- **Health Sciences and** Biological Sciences; and Agricultural Sciences.
- **Multidisciplinary and** Agricultural Sciences.
- **Engineering and** Agricultural Sciences.
- **Biological Sciences and** Agricultural Sciences.

For Q7c, see below:

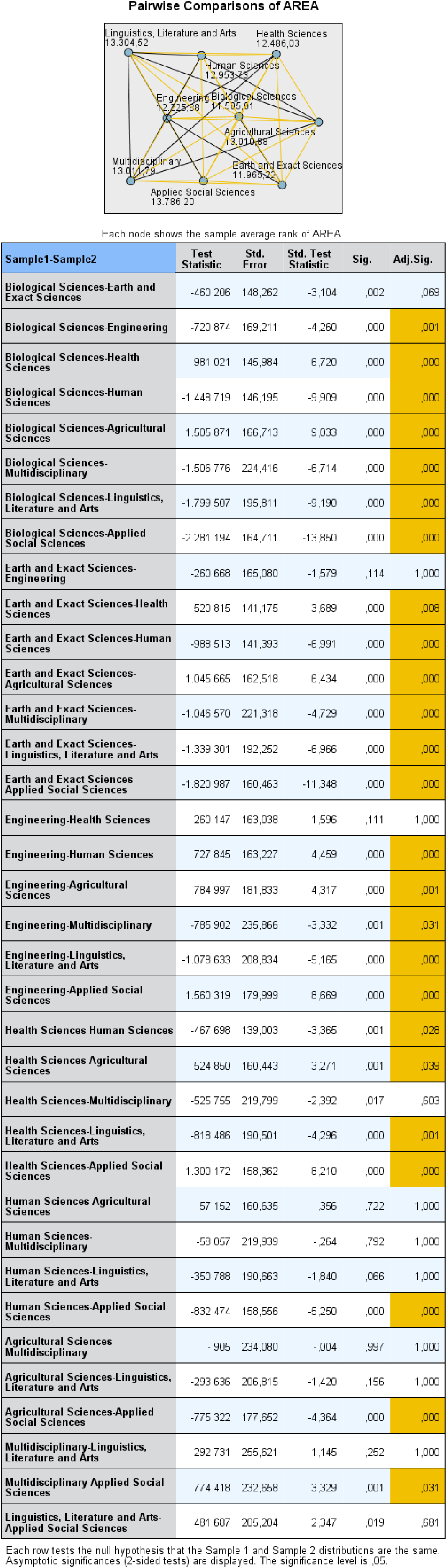

For Q7c, significant differences in response patterns were observed between the following fields:

- **Biological Sciences and** Engineering; and Health Sciences; and Human Sciences; and Agricultural Sciences; and Multidisciplinary; and Linguistics, Literature and Arts; and Applied Social Sciences.
- **Earth and Exact Sciences and** Health Sciences; and Human Sciences; and Agricultural Sciences; and Multidisciplinary; and Linguistics, Literature and Arts; and Applied Social Sciences.
- **Engineering and** Human Sciences; and Agricultural Sciences; Multidisciplinary; and Linguistics, Literature and Arts; and Applied Social Sciences.
- **Health Sciences and** Human Sciences; and Agricultural Sciences; and Linguistics, Literature and Arts; and Applied Social Sciences.
- **Human Sciences and** Applied Social Sciences.
- **Agricultural Sciences and** Applied Social Sciences.
- **Multidisciplinary and** Applied Social Sciences.

For Q8a, see below:

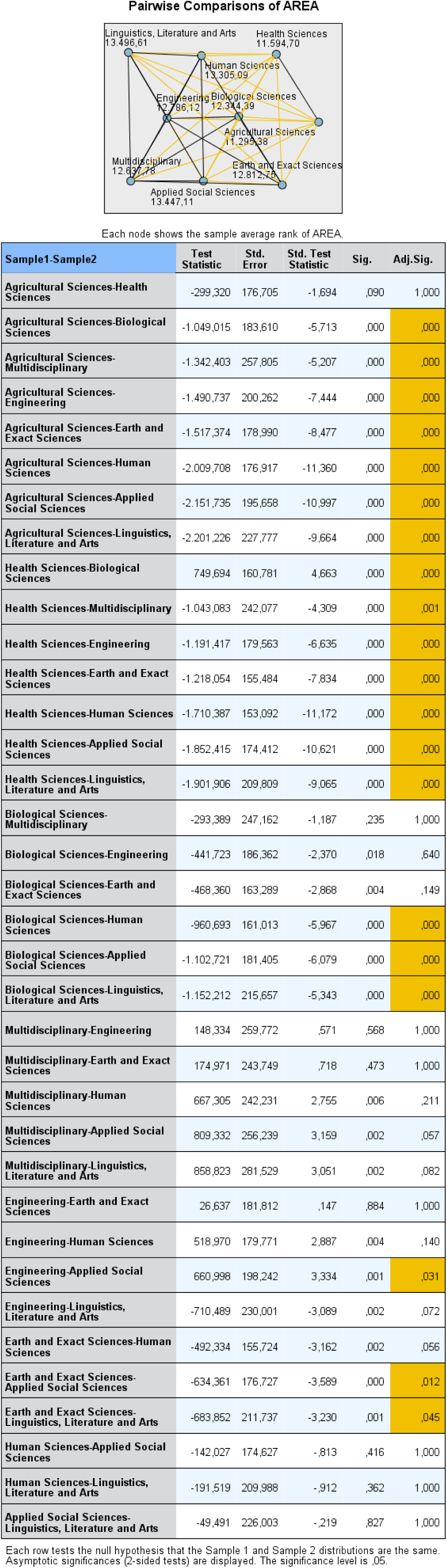

For Q8a, significant differences in response patterns were observed between the following fields:

- **Agricultural Sciences and** Biological Sciences; and Multidisciplinary; and Engineering; and Earth and Exact Sciences; and Human Sciences; and Applied Social Sciences; Linguistics, Literature and Arts.
- **Health Sciences and** Biological Sciences; and Multidisciplinary; and Engineering; and Earth and Exact Sciences; and Human Sciences; and Applied Social Sciences; Linguistics, Literature and Arts.
- **Biological Sciences and** Human Sciences, and Applied Social Sciences; Linguistics, Literature and Arts.
- **Engineering and** Applied Social Sciences;
- **Earth and Exact Sciences and** Applied Social Sciences; and Linguistics, Literature and Arts.

– Note however, that this question is about an issue related to graduate students in the Brazilian context - it would not necessarily involve views or perceptions of respondents about conceptual aspects of plagiarism or the approach to the practice in a particular case.

For Q8b, see below:

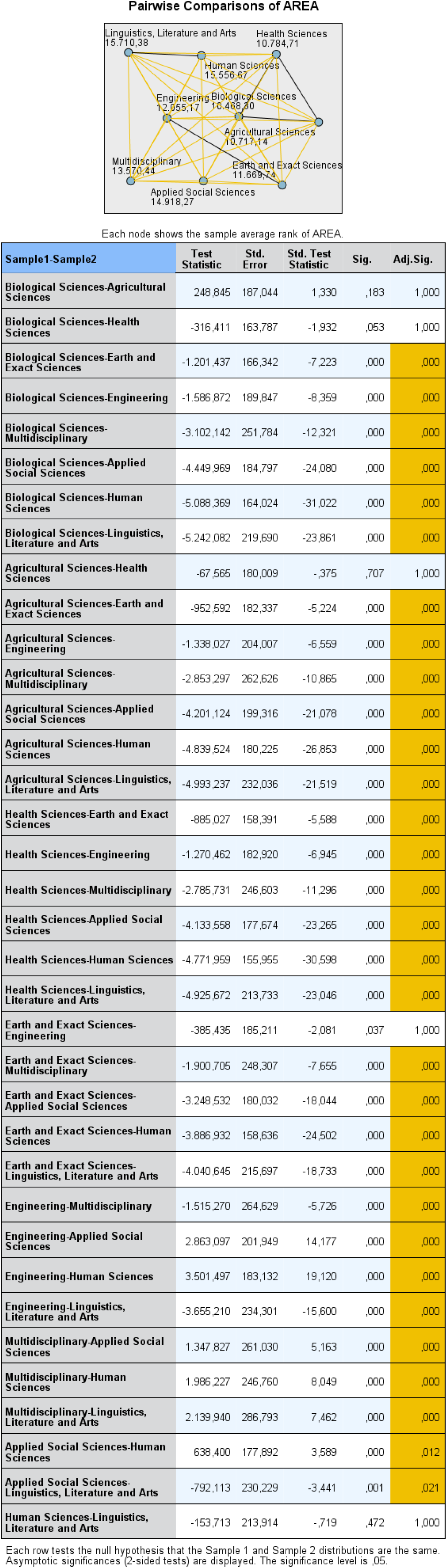

For Q8b, significant differences in response patterns were observed between the following fields:

- **Biological Sciences and** Earth and Exact Sciences; and Engineering; and Multidisciplinary; and Applied Social Sciences; and Human Sciences, Linguistics, Literature and Arts.
- **Agricultural Sciences and** Earth and Exact Sciences; and Engineering; and Multidisciplinary; and Applied Social Sciences; and Human Sciences; and Linguistics, Literature and Arts.
- **Health Sciences and** Earth and Exact Sciences; and Engineering; and Multidisciplinary; and Applied Social Sciences; and Human Sciences; and Linguistics, Literature and Arts.
- **Earth and Exact Sciences and** Multidisciplinary; and Applied Social Sciences; and Human Sciences; and Linguistics, Literature and Arts.
- **Engineering and** Multidisciplinary; and Applied Social Sciences; and Human Sciences; and Linguistics, Literature and Arts.
- **Multidisciplinary and** Applied Social Sciences; and Human Sciences; and Linguistics, Literature and Arts.
- **Applied Social Sciences and** Human Sciences; and Linguistics, Literature and Arts.

– Note however, that this question is about an issue related to graduate students in the Brazilian context - it would not necessarily involve views or perceptions of respondents about conceptual aspects of plagiarism or the approach to the practice in a particular case.

The Hypothesis Test Summary for **Questions in Section III** is as follows:

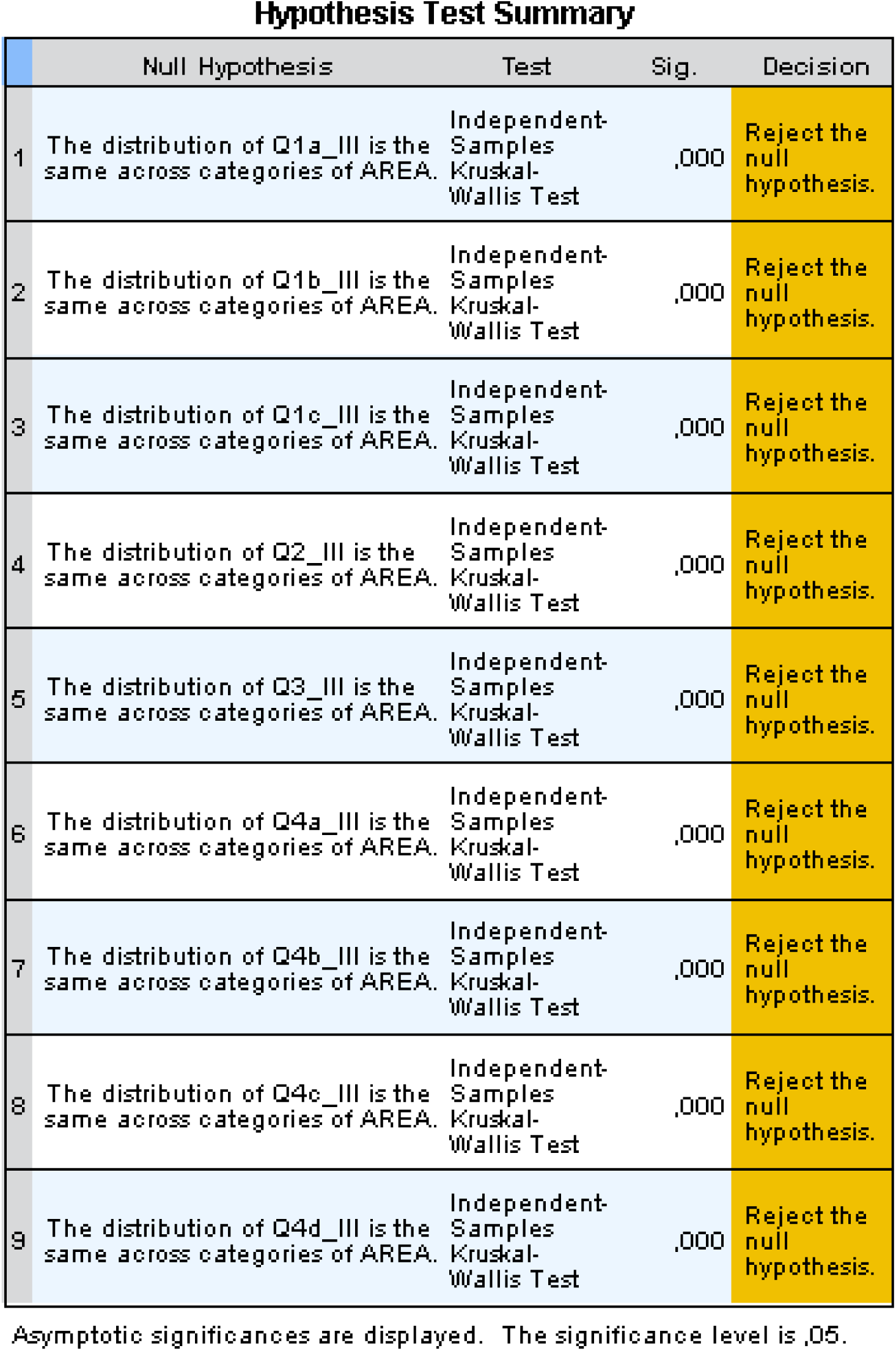

**The differences in response patterns for each question in Section III are listed below:**

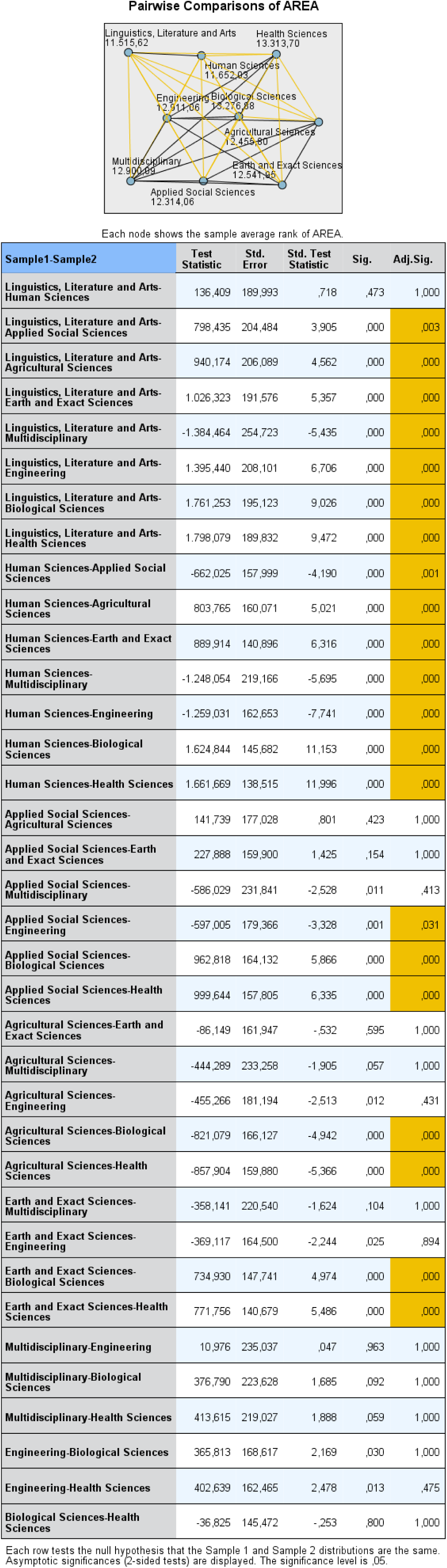

For Q1a, significant differences in response patterns were observed between the following fields:

- **Linguistics, Literature and Arts and** Applied Social Sciences; and Agricultural Sciences; and Earth and Exact Sciences; and Multidisciplinary; and Engineering; and Biological Sciences; and Health Sciences.
- **Human Sciences and** Applied Social Sciences; and Agricultural Sciences; and Earth and Exact Sciences; and Multidisciplinary; and Engineering; and Biological Sciences; and Health Sciences.
- **Applied Social Sciences and** Engineering; and Biological Sciences; and Health Sciences.
- **Agricultural Sciences and** Biological Sciences; and Health Sciences.
- **Earth and Exact Sciences and** Biological Sciences; and Health Sciences.

– Note however, that for Q1a, Q1b, and Q1c, **Section III**, the question is about an issue related to graduate students in the Brazilian context - it would not necessarily involve views or perceptions of respondents about conceptual aspects of plagiarism or the approach to the practice in a particular case.

For Q1b, see below:

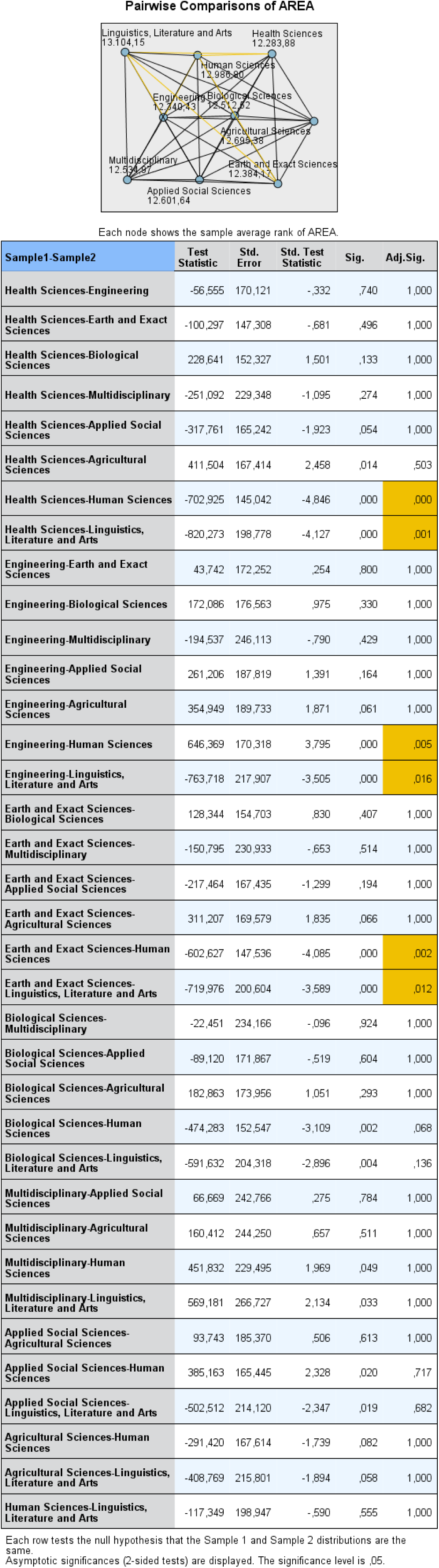

For Q1b, significant differences in response patterns were observed for the following fields:

- **Health Sciences and** Human Sciences; and Linguistics, Literature and Arts.
- **Engineering and** Human Sciences; and Linguistics, Literature and Arts.
- **Earth and Exact Sciences and** Human Sciences; and Linguistics, Literature and Arts.

For Q1c, see below:

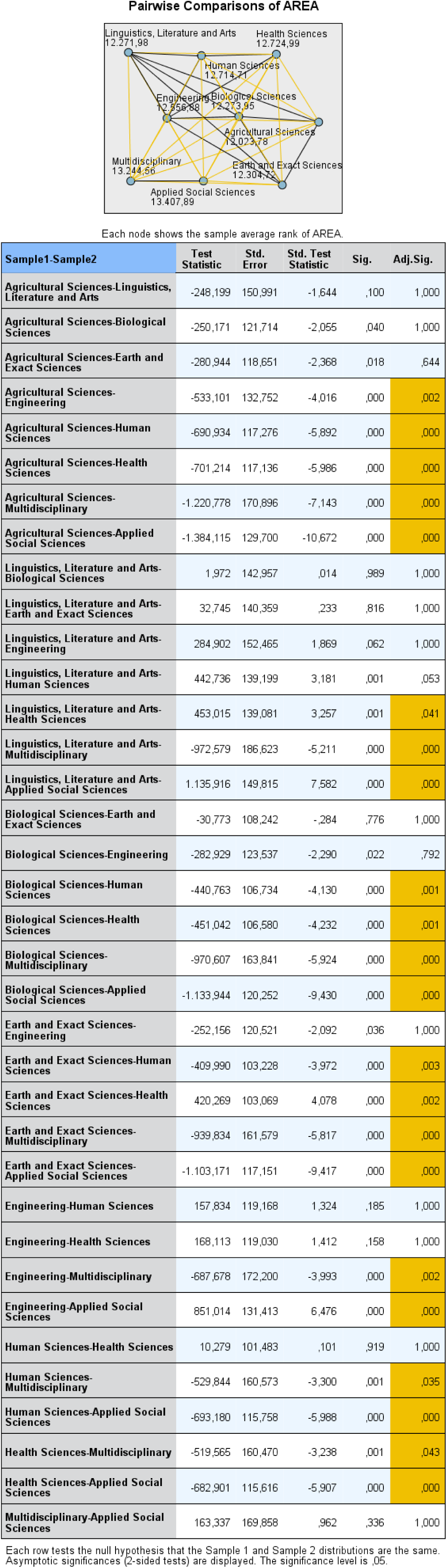

For Q1c, significant differences in response patterns were observed between the following fields:

- **Agricultural Sciences and** Engineering; and Human Sciences, and Health Sciences; and Multidisciplinary; and Applied Social Sciences.
- **Linguistics, Literature and Arts and** Health Sciences; and Multidisciplinary; and Applied Social Sciences.
- **Biological Sciences and** Human Sciences; and Health Sciences; and Multidisciplinary; and Applied Social Sciences.
- **Earth and Exact Sciences and Literature and Arts and** Human Sciences; and Health Sciences; and Multidisciplinary; and Applied Social Sciences.
- **Engineering and** Multidisciplinary; and Applied Social Sciences.
- **Human Sciences and** Multidisciplinary; and Applied Social Sciences.
- **Health Sciences and** Multidisciplinary; and Applied Social Sciences.

For Q2, see below:

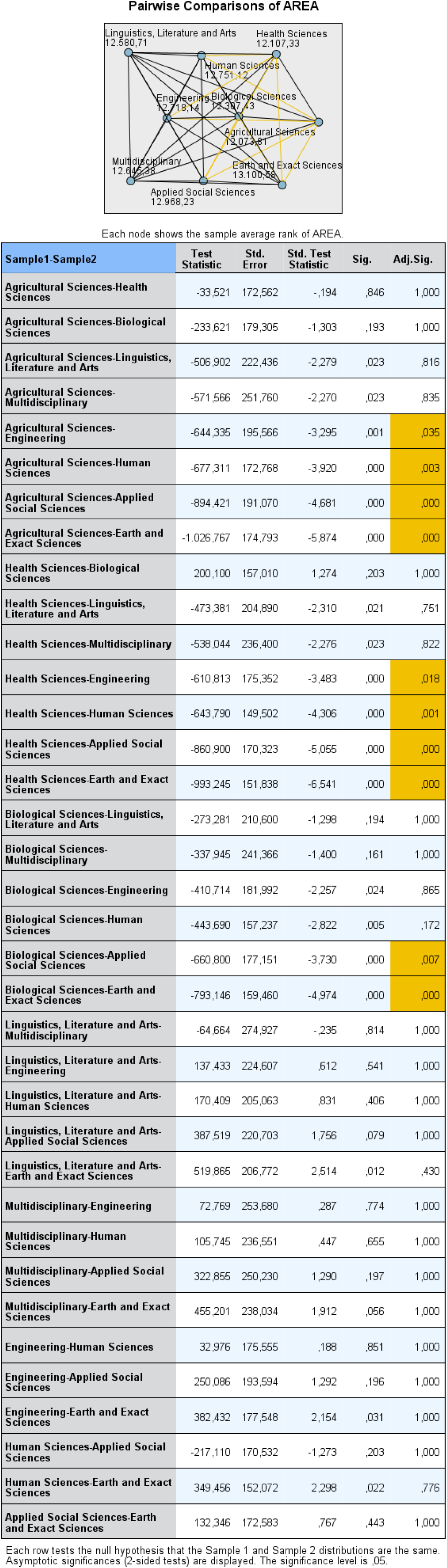

For Q2, significant differences in response patterns were observed between the following fields:

- **Agricultural Sciences and** Engineering; and Human Sciences; and Applied Social Sciences; and Earth and Exact Sciences.
- **Health Sciences and** Engineering, Human Sciences; and Applied Social Sciences. and Earth and Exact Sciences.
- **Biological Sciences and** Applied Social Sciences; and Earth and Exact Sciences.

For Q3, see below:

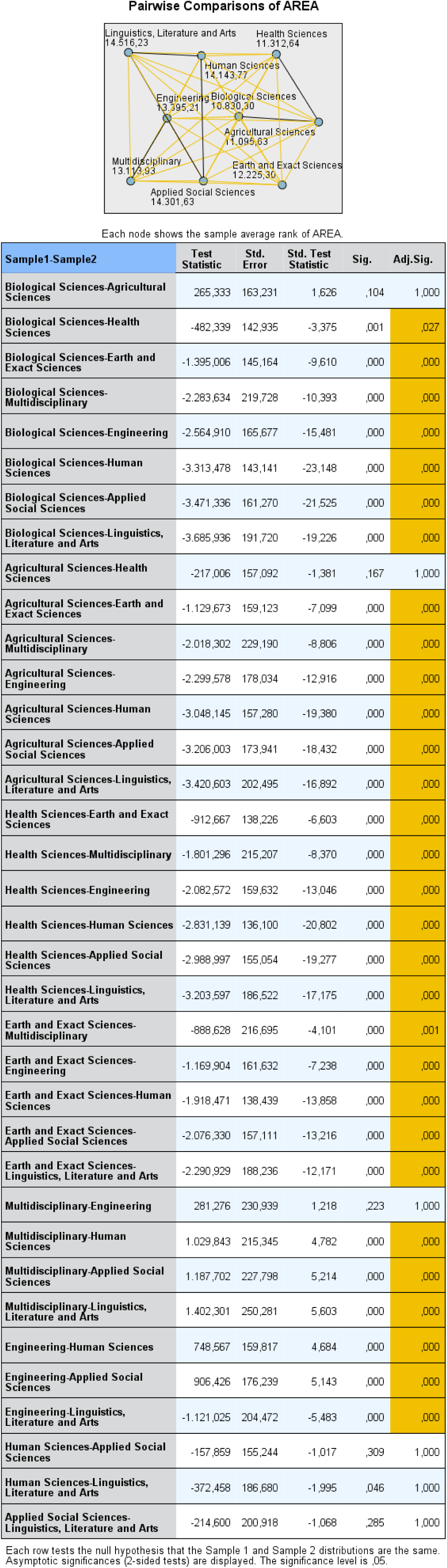

For Q3, significant differences in response patterns were observed between the following fields:

- **Biological Sciences and** Health Sciences; and Earth and Exact Sciences, and Multidisciplinary; and Engineering; and Human Sciences; Applied Social Sciences; Linguistics, Literature and Arts.
- **Agricultural Science**s **and** Earth and Exact Sciences; and Multidisciplinary; and Engineering; and Human Sciences; and Applied Social Sciences; and Linguistics, Literature and Arts.
- **Health Sciences and** Earth and Exact Sciences; an, Multidisciplinary; and Engineering; and Human Science; and Applied Social Sciences; and Linguistics, Literature and Arts.
- **Earth and Exact Sciences and** Multidisciplinary; and Engineering, and Human Sciences; Applied Social Sciences; and Linguistics, Literature and Arts.
- **Multidisciplinary and** Human Sciences; and Applied Social Sciences; and Linguistics, Literature and Arts.
- **Engineering and** Human Sciences; and Applied Social Sciences; and Linguistics, Literature and Arts.

For Q4a, see below:

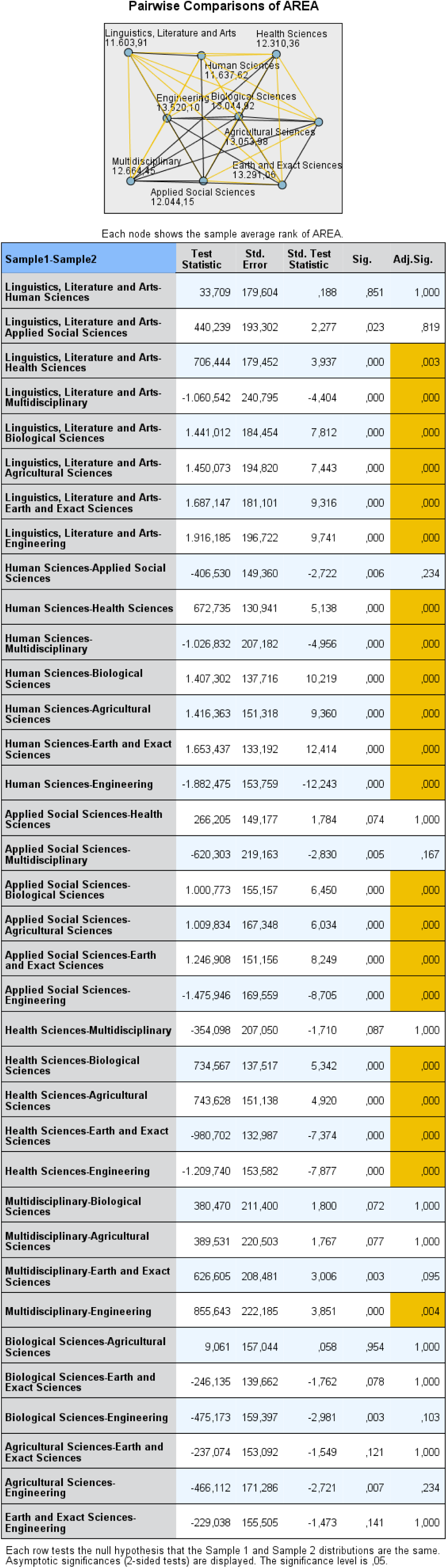

For Q4a, significant differences in response patterns were observed between the following fields:

- **Linguistics, Literature and Arts and** Health Sciences; and Multidisciplinary; and Biological Sciences; and Agricultural Sciences; and Earth and Exact Sciences; and Engineering.
- **Human Sciences and** Health Sciences; and Multidisciplinary; and Biological Sciences; and Agricultural Sciences; and Earth and Exact Sciences; and Engineering.
- **Applied Social Sciences and** Biological Sciences; and Agricultural Sciences; and Earth and Exact Sciences; and Engineering.
- **Health Sciences and** Biological Sciences; and Agricultural Sciences; and Earth and Exact Sciences; and Engineering.
- **Multidisciplinary and** Engineering.

For Q4b, see below:

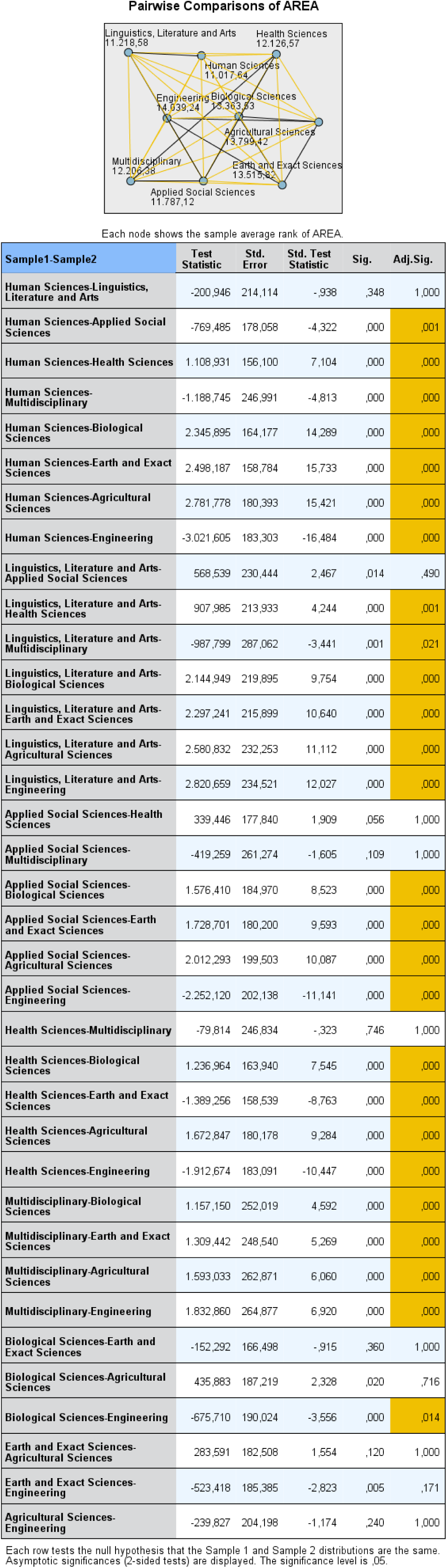

For Q4b, significant differences in response patterns were observed between the following fields:

- **Human Sciences and** Applied Social Sciences; and Health Sciences; and Multidisciplinary; and Biological Sciences, and Earth and Exact Sciences; and Agricultural Sciences; and Engineering.
- **Linguistics, Literature and Arts and** Health Sciences; and Multidisciplinary; and Biological Sciences; and Earth and Exact Sciences; and Agricultural Sciences and Engineering.
- **Applied Social Sciences and** Biological Sciences; and Earth and Exact Sciences, and Agricultural Sciences; and Engineering.
- **Health Sciences and** Biological Sciences; and Earth and Exact Sciences, and Agricultural Sciences; and Engineering.
- **Multidisciplinary and** Biological Sciences; and Earth and Exact Sciences, and Agricultural Sciences; and Engineering.
- **Biological Sciences and** Engineering.

For Q4c, see below:

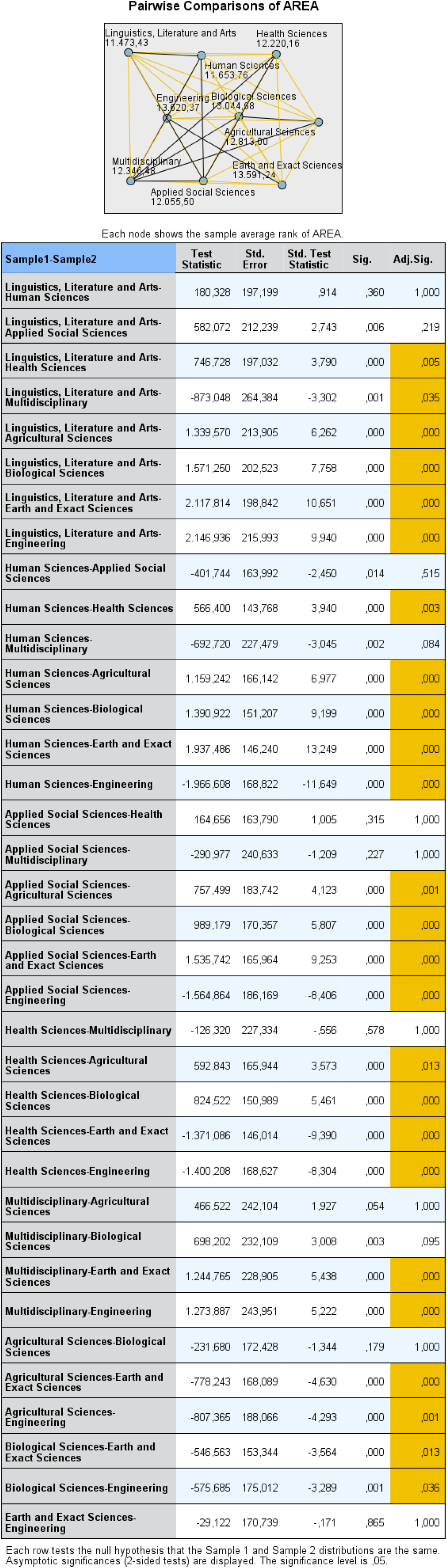

For Q4c, significant differences in response patterns were observed between the following fields:

- **Linguistics, Literature and** Arts and Health Sciences; and Multidisciplinary; and Agricultural Sciences; and Biological Sciences; and Earth and Exact Sciences; and Engineering.
- **Human Sciences and** Health Sciences; Agricultural Sciences; and Biological Sciences; and Earth and Exact Sciences and Engineering.
- **Applied Social Sciences and** Agricultural Sciences; and Biological Sciences; and Earth and Exact Sciences; and Engineering.
- **Health Sciences and** Agricultural Sciences; and Biological Sciences; and Earth and Exact Sciences; and Engineering.
- **Multidisciplinary and** Earth and Exact Sciences; and Engineering.
- **Agricultural Sciences and** Earth and Exact Sciences; and Engineering.
- **Biological Sciences and** Earth and Exact Sciences; and Engineering.

For Q4d, see below:

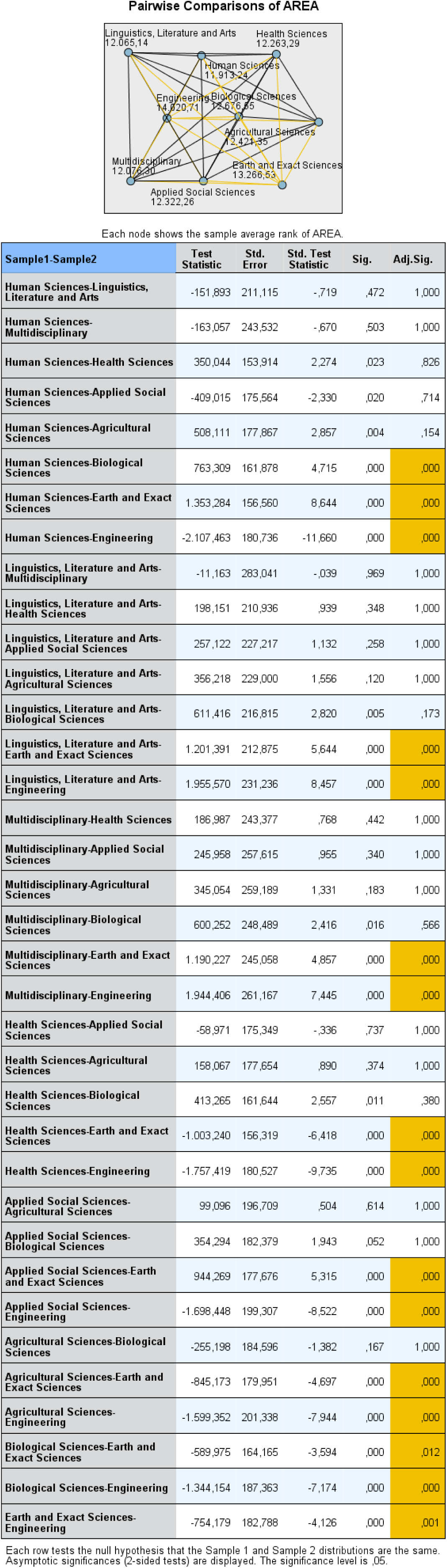

For Q4d, significant differences in response patterns were observed between the following fields:

- **Human Sciences and** Biological Sciences; and Earth and Exact Sciences; and Engineering.
- **Linguistics, Literature and Arts** and Earth and Exact Sciences; and Engineering.
- **Multidisciplinary and** Earth and Exact Sciences; and Engineering.
- **Health Sciences and** Earth and Exact Sciences; and Engineering.
- **Applied Social Sciences and** Earth and Exact Sciences; and Engineering.
- **Agricultural Sciences and** Earth and Exact Sciences; and Engineering.
- **Biological Sciences and** Earth and Exact Sciences; and Engineering.
- **Earth and Exact Sciences and** Engineering.

## Letter of Invitation by the Principal Investigator and Confidentiality Statement

Welcome!

This questionnaire is divided into five steps:

General Information (**Section I**)
Questions about plagiarism (**Section II**)
Questions regarding self-plagiarism (**Section III**)
Questions about redundancy (**Section IV**)
Your comments and suggestions (**Section V**)

Conditions: responses and information recorded in the questionnaire

a. will be used only for the goals described in the invitation letter
b. will be taken as personal opinions
c. will not be considered general (institutional) opinions

To start the questionnaire, click on the tab **START**

This action will tell us that you agree with the conditions.

To start a new section, you should complete the previous one.

START

### In this first step of the questionnaire (Section I), our goal is to collect information about you. This information will help us to understand the responses to other sections of this questionnaire

DATE OF BIRTH
SEX
**Female**
**Male**
WHAT IS YOUR LAST DEGREE EARNED?
**Bachelors**
**Specialization**
**Masters**
**PhD**
MAIN AREA OF ACADEMIC EXPERTISE
**Exact and Earth Sciences**
**Biological Sciences**
**Engineering**
**Health Sciences**
**Agricultural Sciences**
**Applied Social Sciences**
**Human Sciences**
**Language, Literature and Arts**
**Multidisciplinary**

IF YOU HAVE A PhD, IN WHICH STATE WAS YOUR DOCTORATE AWARDED?

[ALL BRAZILIAN STATES LISTED HERE]

IF NOT BRAZIL, PLEASE INDICATE THE COUNTRY IN WHICH YOU OBTAINED YOUR DOCTORAL TRAINING:

[LIST OF COUNTRIES]

YEAR OF PHD COMPLETION

DID YOU TAKE A POST-DOC?

IF NOT BRAZIL, PLEASE INDICATE THE COUNTRY IN WHICH YOU OBTAINED YOUR POST-DOCTORAL TRAINING

[LIST OF COUNTRIES]

YOUR INSTITUTION OF AFFILIATION IN 2013

**Public**
**Private**
**Both**

IN WHAT STATE IS YOUR INSTITUTION LOCATED

[ALL BRAZILIAN STATES LISTED HERE]

POSITION AT THE INSTITUTION WHERE YOU WORKED IN 2013:

**Substitute Professor**
**Assistant OR Associate Professor**
**Full Professor**
**Visiting Professor**
**Emeritus Professor**
**Retired Professor**
**Researcher (non-teaching activity)**
**\* This last option includes post-docs who are not professors**

NEXT

### For Section II, our goal is to obtain information about your perceptions of plagiarism and related issues

1. The definition of misconduct established in 2000 by the U.S. Office of Science and Technology Policy (OSTP) is as follows: “Research misconduct is defined as fabrication, falsification, or plagiarism in proposing, performing, or reviewing research, or in reporting research results.” Thus, just as fabrication and falsification are research misconduct, so is plagiarism. Do you agree?

Agree
Partially agree
I don’t know
Partially disagree
Disagree

2. The definition of academic plagiarism by OSTP, embraced by much of the international academic community, is as follows: “Appropriation of another person’s ideas, processes, results, or words without giving appropriate credit”.

2a. Do you consider this definition clear?

Yes
Yes, partially
I don’t know
No, the definition is greatly simplified
No, the definition is confusing
No

2b. Additional Comments

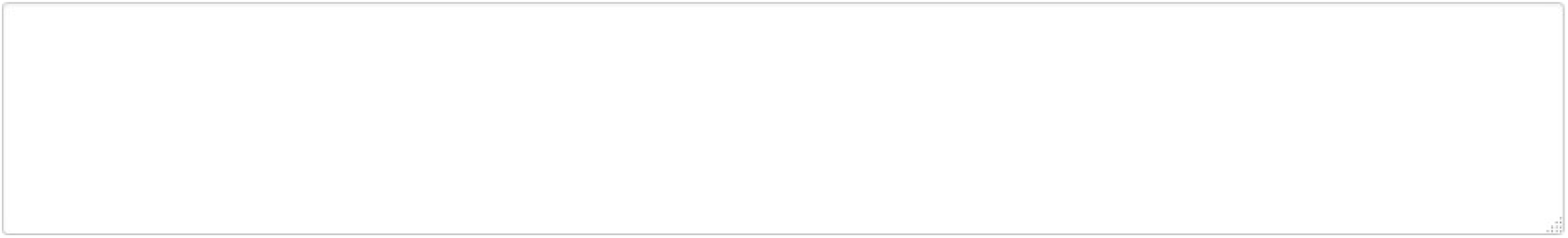

3. Do you agree with the above definition of plagiarism?

Agree
Partially agree
I don’t know
Partially disagree
Disagree

4. How do you see plagiarism in science?

Nothing to worry about in the scientific community
An error, but not research misconduct
An unethical practice, but not research misconduct
Research misconduct
Research misconduct, except for textual plagiarism
Research misconduct, except for plagiarism of ideas

5. Recent surveys indicate an increase in plagiarism in scientific publications. Many of the cases have led to “retractions” of scientific papers (cancellation of publications). In this context, in 2010, *Nature Reviews Genetics* (NRG) retracted a review article for textual plagiarism [*Nature Reviews Genetics* 11:308(2010]. The plagiarism involved a single paragraph that had been paraphrased from an article submitted to *Plant Science*. The author of the NRG review was a referee for the *Plant Science* paper but she failed to cite it when she wrote the NRG review. In the retraction notice the NRG editors stated that the misappropriated paragraph was plagiarized and that the author of the NRG review had presented the ideas and hypotheses found in the original paragraph as if they were her own.

5a. Do you agree that textual plagiarism justifies a retraction?

Agree
Partially agree
I don’t know
Partially disagree
Disagree

5b. Do you agree with the retraction in this case?

Agree
Partially agree
I don’t know
Partially disagree
Disagree

5c. Additional Comments

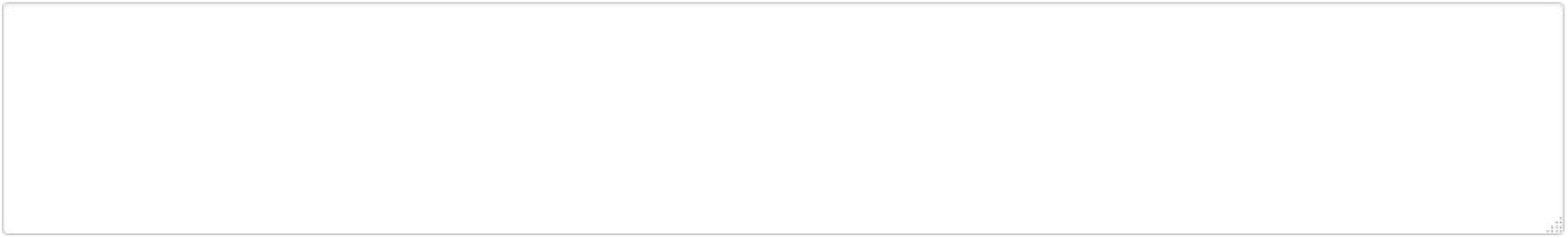

6. Plagiarism in graduate school has become a concern in most universities in the world, including Brazil. In October 2010, a document from the Brazilian Bar Association (OAB) expressed concern on this topic in the academic context of our country.

Have you read that document?

Yes
Yes, partially
I do not remember
No, but I knew of the existence of this document
No, I did not know of the existence of this document

7. Have you ever encountered a case of plagiarism (partial or total) by a graduate student (not necessarily from your program) in the last four years?

7a. in reviewing a Master’s thesis

No
Yes, one case
Yes, two cases
Yes, three cases
Yes, more than three cases
Yes, many cases

7b. in reviewing a PhD thesis

No
Yes, one case
Yes, two cases
Yes, three cases
Yes, four cases
Yes, many cases

7c. in reviewing a scientific paper

No
Yes, one case
Yes, two cases
Yes, three cases
Yes, four cases
Yes, many cases

8. In the context of plagiarism, it is worth noting that in Brazil:

8a. Graduate students are not familiar with the international concept of academic plagiarism. Agree

Agree
Partially agree
I don’t know
Partially disagree
Disagree

8b. Graduate students tend to engage in textual plagiarism in scientific articles in English because they are not fluent in English.

Agree
Agree
Partially agree
I don’t know
Partially disagree
Disagree

PREVIOUS NEXT

### In Section III of the questionnaire, our goal is to address some issues that have proven relevant to the discussion of self-plagiarism in scientific publications

1. Consider the following information: “Probably the most widely used program to spot plagiarism in scientific publishing is Crosscheck [link], Iaunched in June 2008 by CrossRef. A total of 119 publishers (nearly 50,000 journals) subscribe to the plagiarism detection program, including Elsevier, Wiley-Blackwell, and Springer…” (When is self-plagiarism ok? The Scientist, Sep 2010)

1a. Have you ever heard of such software used by scientific publishers?

Yes
No

1b. Do you consider the use of such software an effective measure to identify scientific plagiarism?

Yes
I don’t know
No

1c. Do you use some type of plagiarism detection software to evaluate the originality of your own manuscript prior to submitting it to a journal?

Yes
No

2. Cases of self-plagiarism in science have been claimed using Crosscheck. However, there is little consensus as to how much material reused by an author, borrowing from his own publication, would be self-plagiarism An author recently accused of self-plagiarizing from one of his previously published articles claimed: “I cannot plagiarize myself; those words are mine.” (When is self-plagiarism ok? The Scientist, Sep 2010). Do you agree with this view?

Agree
Partially agree
I don’t know
Partially disagree
Disagree

2a. Additional Comments

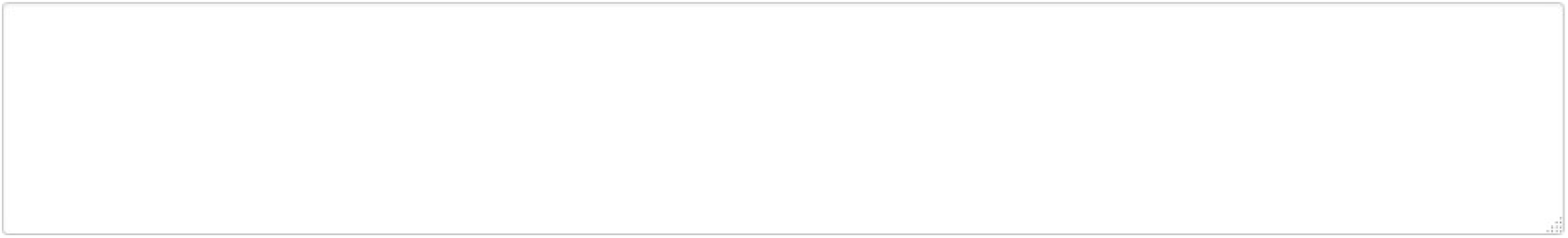

3. Consider the following case: a group of authors submitted a manuscript to a high impact, English language journal that had been previously published in a low impact, German-language journal, in the belief that this publication in English would fulfill the need for higher visibility. The manuscript, which was accepted and published, contained no reference to the publication in German. Although written in different languages, the articles were identical, and both were later retracted after a complaint from a colleague. Do you agree with the retraction?

Agree
Partially agree
I don’t know
Partially disagree
Disagree

3a. Additional Comments

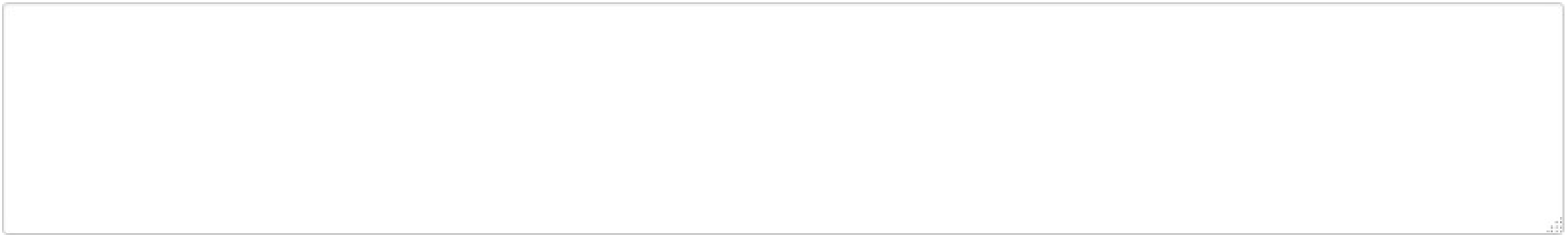

4a. The originality of the results in a research paper should be questioned if the author of that paper **copied entire paragraphs without citation from others’ previously published papers.**

Agree
Partially agree
I don’t know
Partially disagree
Disagree

4b. The originality of the results in a research paper should be questioned if the author of that paper **copied entire paragraphs from others’ previously published papers, citing these sources but without enclosing the copied text in quotation marks.**

Agree
Partially agree
I don’t know
Partially disagree
Disagree

4c. The originality of the results in a research paper should be questioned if the author of that paper correctly paraphrased **entire paragraphs from others’ previously published papers but without citing the original sources.**

Agree
Partially agree
I don’t know
Partially disagree
Disagree

4d. The originality of the results in a research paper should be questioned if the author of that paper copied **entire paragraphs without citation from other previously published papers, and that same person was a co-author of all the publications involved.**

Agree
Partially agree
I don’t know
Partially disagree
Disagree

4e. Additional Comments

PREVIOUS NEXT

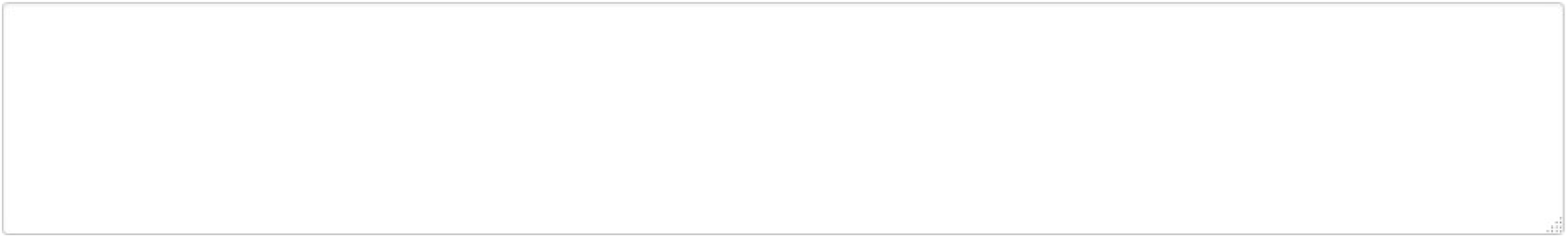

### In Section IV of the questionnaire, our goal is to assess how you view redundancy in scientific publications

1. Consider the following description of redundant publications:

“Redundant, duplicate, or repetitive publications occur when there is representation of 2 or more studies, data sets, or publications in either electronic or print media. The publications may overlap partially or completely, such that a similar portion, major component(s), or complete representation of a previously/simultaneously or future published study is duplicated. These publications may share the same, similar, or overlapping data, hypotheses, discussion, methods, results, and/or conclusions.” (JMPT, 2006, 29, 7: 505-509)

1a. “Editors and authors were in consensus that redundant publications occur because authors feel pressure to publish.” (J Med Ethics, 29:109-114). Do you share this view?

Agree
Partially agree
I don’t know
Partially disagree
Disagree

1b. Additional Comments

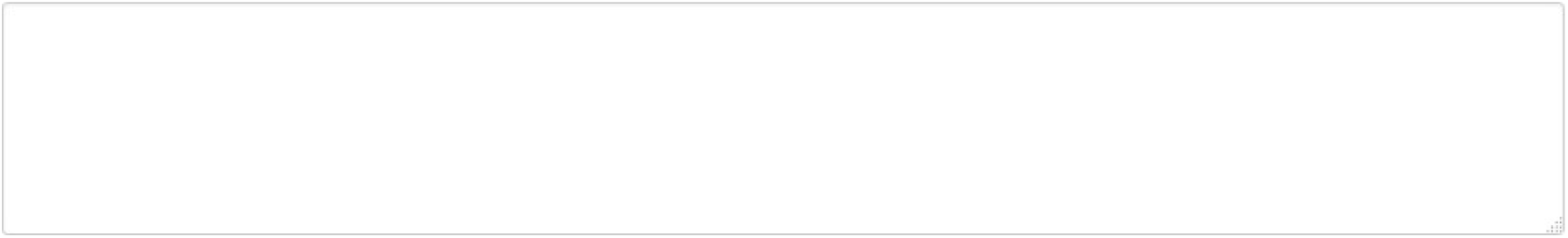

1c. Redundant publications occur because academic leaders do not publicly condemn the practice, because authors do not understand how redundant reporting distorts the aggregation of data… and because authors want to disseminate their research as widely as possible”. (J Med Ethics 2003,29:109-114). Do you agree with this view?

Agree
Partially agree
I don’t know
Partially disagree
Disagree

1d. Additional Comments

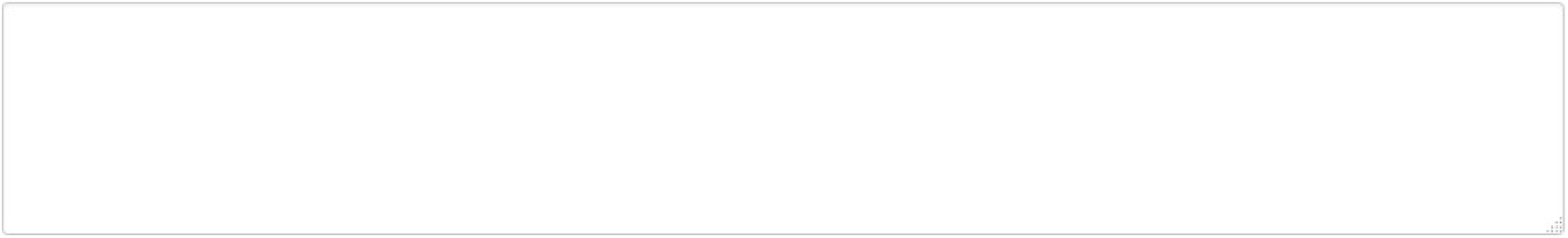

1e. “… Authors should sign statements for journals attesting that their manuscript does not overlap substantially with other of their articles.” (J Med Ethics 2003; 29:109-114)

Do you agree that this should be formalized in writing at the time of submission?

Agree
Partially agree
I don’t know
Partially disagree
Disagree

1f. Additional Comments

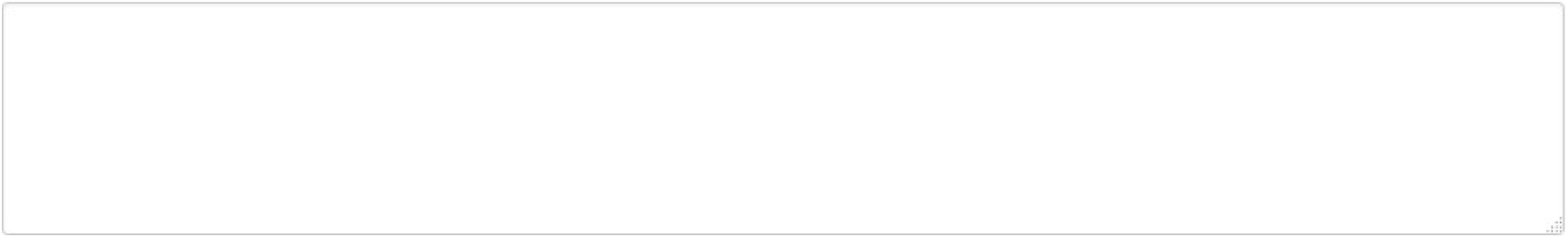

2. In a study published in 2005 (Nature 435:737-738) which surveyed 3,247 U.S. researchers about certain practices that may be considered unethical in science, about 4.7% of respondents admitted to having posted the same result in two or more publications in the previous three years.

2a. Do you consider this an unethical practice?

Yes
I don’t know
No

2b. What is your perception of this practice (publishing the same result in two or more publications) in Brazil?

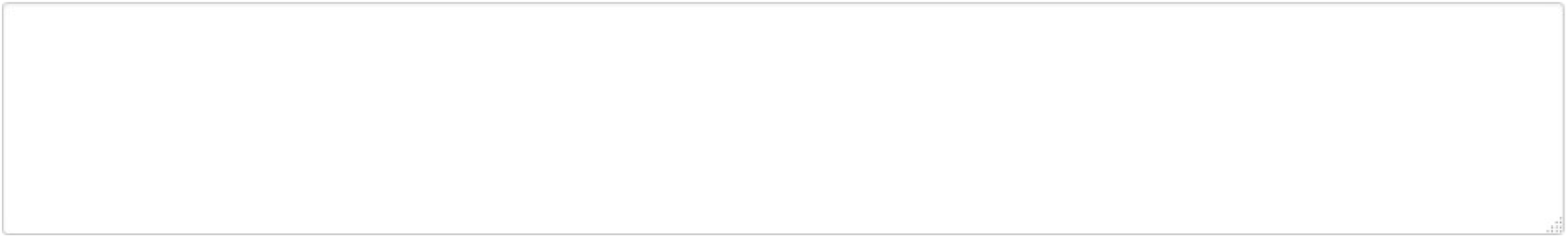

At this last stage, you can provide your comments and suggestions on measures that, in your view, could contribute to eliminating the production of academic work that contains plagiarized or redundant material by Brazilian researchers.

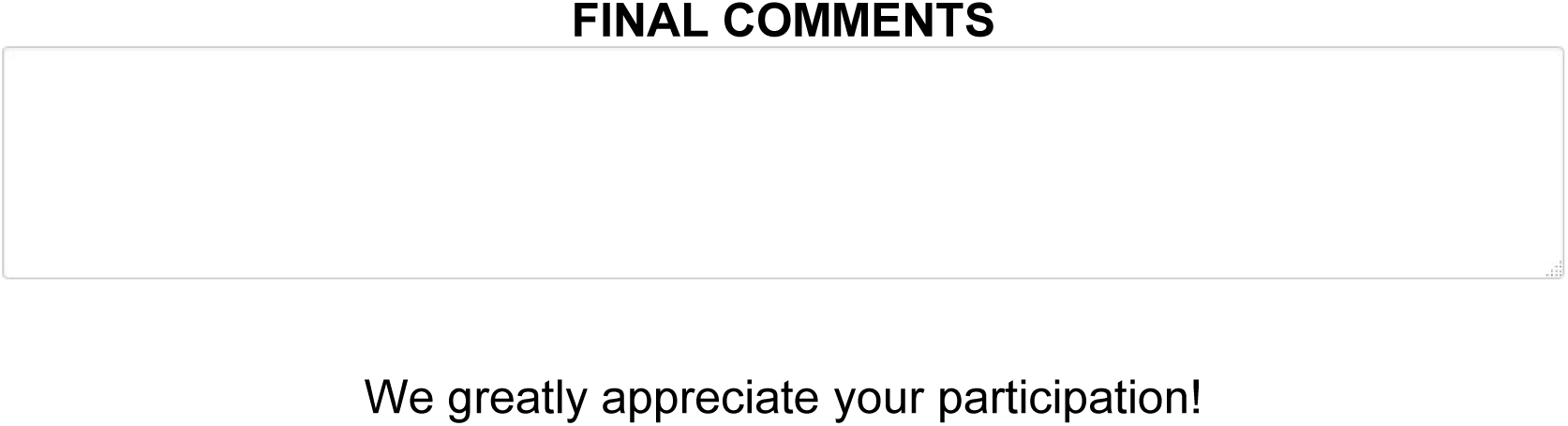

**-- References and list of collaborators added**

